# Binge feeding-induced Olfactory Cortex Suppression Reduces Satiation

**DOI:** 10.1101/2023.10.17.562714

**Authors:** Hung Lo, Malinda L.S. Tantirigama, Anke Schoenherr, Laura Moreno-Velasquez, Lukas Faiss, Benjamin R. Rost, Matthew E. Larkum, Benjamin Judkewitz, Katharina Stumpenhorst, Marion Rivalan, York Winter, Dietmar Schmitz, Friedrich W. Johenning

## Abstract

Binge eating commonly leads to overeating, but the exact mechanism is unclear. While it is known that experiencing flavor contributes to satiety, the interactions between flavor, feeding rate, and food intake remain unknown. Here, we demonstrate a novel feeding rate-dependent feedback loop between olfactory flavor representation in the anterior olfactory (piriform) cortex (aPC) and food intake. Using miniscopes for *in vivo* calcium imaging in freely foraging mice, we identified specific excitatory neuronal responses to food and water during slow feeding. Switching to binge feeding transformed these specific responses into unspecific global suppression of neuronal activity. Food consumption was predicted by the degree of suppression of neuronal activity in the aPC during binge feeding. Also, food deprivation enhanced neuronal activity suppression. We confirmed the hypothesis that aPC suppression promotes food intake with closed-loop optogenetics experiments. Together, we show that olfactory sensory representation in the aPC reciprocally interacts with consummatory behavior to enhance food intake.

## INTRODUCTION

Eating rapidly in a short period, commonly known as binge eating, reduces satiation, the process leading to satiety. Therefore, eating proceeds beyond homeostatic needs (Scisco et al. 2011; Scisco et al. 2011; Andrade, Greene, and Melanson 2008; Bolhuis et al. 2013; Teo, Dam, and Forde 2020; Hurst and Fukuda 2018; Bolhuis et al. 2014). Satiation is an important feedback signal that reduces food consumption upon food intake. The soup paradox illustrates the intimate relationship between feeding rate and satiation: Energy-dense liquids, like apple juice, offer less long-term satiety compared to their isocaloric solid counterparts, such as apples. Interestingly, however, when the rate of liquid food intake is slowed down, for example, with spoon feeding, its ability to satisfy hunger matches that of solid foods, influencing 24-hour food intake (Mattes 2005). The canonical explanation for the reduction of satiation by binge feeding in comparison to slow feeding is based on the delayed transfer of homeostatic signals from the gastrointestinal tract to the brain (Slyper 2021; Samakidou et al. 2023; Grove et al. 2022). While visceral satiation based on ingestion and absorption is undisputed, there is also sensory satiation mediated by flavor perception (Chen et al. 2015; Mandelblat-Cerf et al. 2015; Cecil, Francis, and Read 1999; Betley et al. 2015). However, no study has investigated whether alterations in flavor representation during binge feeding can reduce sensory satiation. Such a feedback loop between feeding behavior, appetite, and flavor representation would support the emerging concept that reciprocal interactions between action and perception shape behavior (Buzsáki 2019).

Flavor is a multisensory phenomenon involving multiple interconnected primary sensory areas including the primary olfactory cortex or piriform cortex (PC) representing smell, the gustatory cortex (GC) representing taste, and, to some degree, other sensory cortices for tactile and visual aspects of food (Small 2012; Elliott and Maier 2020). The PC generally represents odor identity in a concentration-invariant population code of activated neurons (Blazing and Franks 2020). Random activation of neuronal subpopulations in the PC can determine conditioned appetitive and aversive behavioral responses (Choi et al. 2011), and the PC population code does not contribute to the signaling of odor valence (Wang et al. 2020). If the PC’s role in flavor representation during feeding were limited to identification without affecting specific behavioral outcomes, we would expect a stable representation of flavors in the neuronal response patterns to slow and binge feeding.

## RESULTS

### Feeding rate modulates flavor representation in the anterior piriform cortex

To examine how feeding rate affects sensory representation, we established a feeding paradigm combined with calcium (Ca^2+^) imaging in freely moving mice. We built a liquid food delivery system that feeds mice at two different rates to experimentally induce slow or binge feeding. Mice could voluntarily lick from a lick spout to trigger the delivery of a droplet (∼1.8 µL) of liquid food (Ensure, artificial energy-dense flavored nutrient solution) or water. Over 30 min, mice had access to the spout for a total of 14 minutes, with a pseudorandom order of 2 min long slow and fast feeding periods. To control the feeding rate, we implemented different refractory periods for the delivery pump: 4 seconds for the slow feeding mode and 0.4 seconds for the binge feeding mode (Fig. 1A). Mice consumed more food and licked at a higher rate during binge feeding (Fig. S1A-D).

**Figure. 1.**
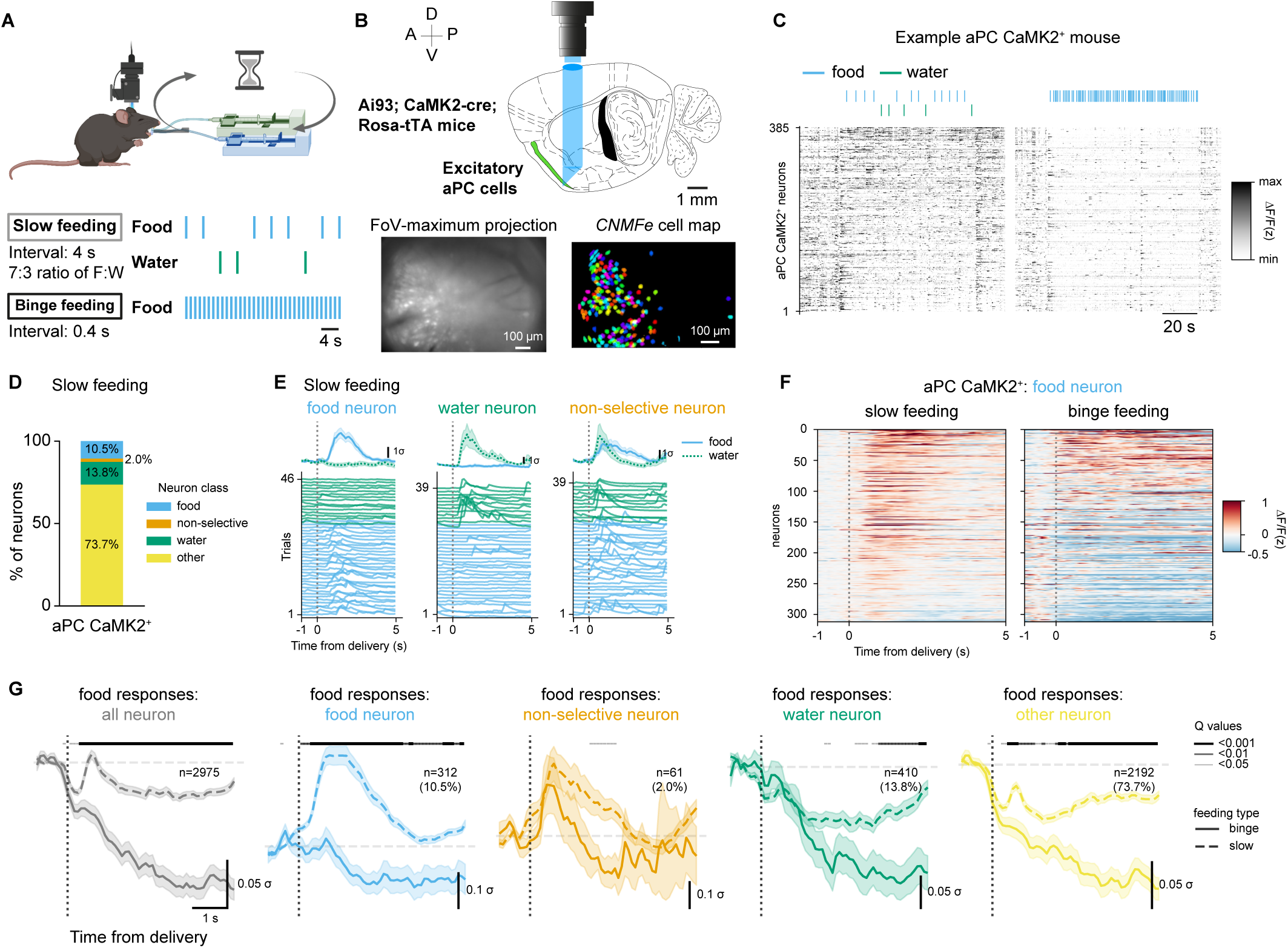
Feeding rate modulates flavor representations in the aPC. (**A**) Miniscope recording and behavioral protocol for slow feeding and binge feeding. (**B**) Top: Schematics of GRIN lens/Prism (blue shade) implantation in the aPC (green structure). Bottom: Field-of-view of miniscope recordings and extracted cell maps by constrained non-negative matrix factorization (*CNMFe*). (**C**) Example Ca^2+^ traces of aPC neurons during slow feeding and binge feeding from one mouse. (**D**) Percentage of aPC neurons activated during slow feeding by food, water, or non-selective consumption vs. non-responding neurons. (n=2975 cells, 481 slow feeding trials, and 241 binge feeding trials in 8 mice). (**E**) Trial-average and single trial responses of example aPC food-activated, water-activated, and non-selective neurons upon slow feeding. (**F**) Trial-averaged responses to food deliveries of individual aPC CaMK2^+^ food-activated neurons upon slow feeding and binge feeding (n=312 cells in 8 mice). (**G**) Trial-average responses of population and subclasses of aPC CaMK2^+^ neurons upon slow and binge feeding (n is the same as in **D**). The dashed vertical line indicates the start of the food delivery. The shaded line above denotes the adjusted P-values (Q-values) of each time point, with different line widths representing different values (from thin to thick: Q <0.05, <0.01, <0.001). For (**E**) and (**G**) data are shown as mean ± s.e.m.

To record feeding-related neuronal activity patterns in the anterior olfactory (piriform) cortex (aPC), we used endomicroscopic lenses (GRIN lenses) attached to prisms combined with a miniscope to image Ca^2+^ transients of excitatory aPC neurons expressing GCaMP6f (Fig. 1B, and S3A-C). We were able to stably record around 140 aPC excitatory neurons per individual imaging session (Fig. S3D, average = 140.40 ± 72.19 neurons).

When switching from slow feeding to binge feeding, we observed a robust suppression of the aPC excitatory neuron population activity (Fig. 1C, Supplementary Video S1). We categorized neurons based on their responses during slow feeding, as this feeding mode provided a larger number of clearly separated individual trials. We interspersed brief water deliveries in the slow feeding mode to differentiate food-specific neurons from non-specific feeding-related neurons. This protocol allowed us to classify neurons pooled over animals and sessions into food-activated (10.5%), water-activated (13.8%), non-selective consumption-activated (neurons responding to both food and water deliveries; 2.0%), and non-responders (73.7%; Fig. 1D, E and S2A, B). Binge feeding-induced aPC suppression was present in food-activated, water-activated, and non-responding neurons but not in non-selective consumption-activated neurons (Fig. 1F, G, S2C-F, statistical analysis of the neuronal activity differences during slow and binge feeding are illustrated with Q-values (unpaired t-test with false discovery rate correction) along the averaged Ca^2+^ traces).

Activity in the PC represents odor identity through a distributed population code and transmits information about odor identity and concentration to downstream targets (Miura, Mainen, and Uchida 2012; Stettler and Axel 2009; Wilson and Sullivan 2011; Bolding and Franks 2017; Berners-Lee et al. 2023; Tantirigama, Huang, and Bekkers 2017). The percentage of cells in the food-specific subclass during slow feeding is comparable to odor-specific populations observed in the aPC (Tantirigama, Huang, and Bekkers 2017; Roland et al. 2017). When odors are presented at different intensities, the distributed population code and the neuronal firing rates remain similar (Bolding and Franks 2017). So, in spite of the higher food volume over time during binge feeding resulting in increased intensity of flavors of the same identity, we did not expect changes in the population code (Fig. 1g, 2E). Thus, our data points to a feeding rate-dependent modulation of flavor representation in the aPC that fundamentally alters the flavor-specific population code.

### Flavor representation in the gustatory cortex is stable across feeding rates

In general in sensory systems, behavior has profound effects on brain activity underlying the representation of sensory inputs (Musall et al. 2019; Steinmetz et al. 2019; Stringer et al. 2019). Therefore, neuronal suppression in the aPC induced by binge feeding could be a more global phenomenon that might be observed in other brain regions involving flavor representation. To test this, we performed Ca^2+^ imaging in the gustatory cortex (GC, granular, and dysgranular insular cortex) using a miniscope and tracked taste representation during slow and binge feeding (Fig. 2A-C, S4A-G). We found little modulation in the general population of GC neurons upon binge feeding compared with slow feeding (Fig. 2D), except for prolonged activation of GC neurons during binge feeding (Fig. 2D). in GC water-activated, non-selective and non-responding neurons, binge feeding of food had minimal levels of modulation compared with the general suppression in aPC neurons. (Fig. 2E, Fig. S5A; estimated effect sizes in each neuron class for aPC and GC neurons). Unlike the global activity suppression of the food-activated neurons in the aPC, food-specific activation of GC neurons was selectively preserved during binge feeding (Fig. 2D, E, and S5B).

**Figure. 2.**
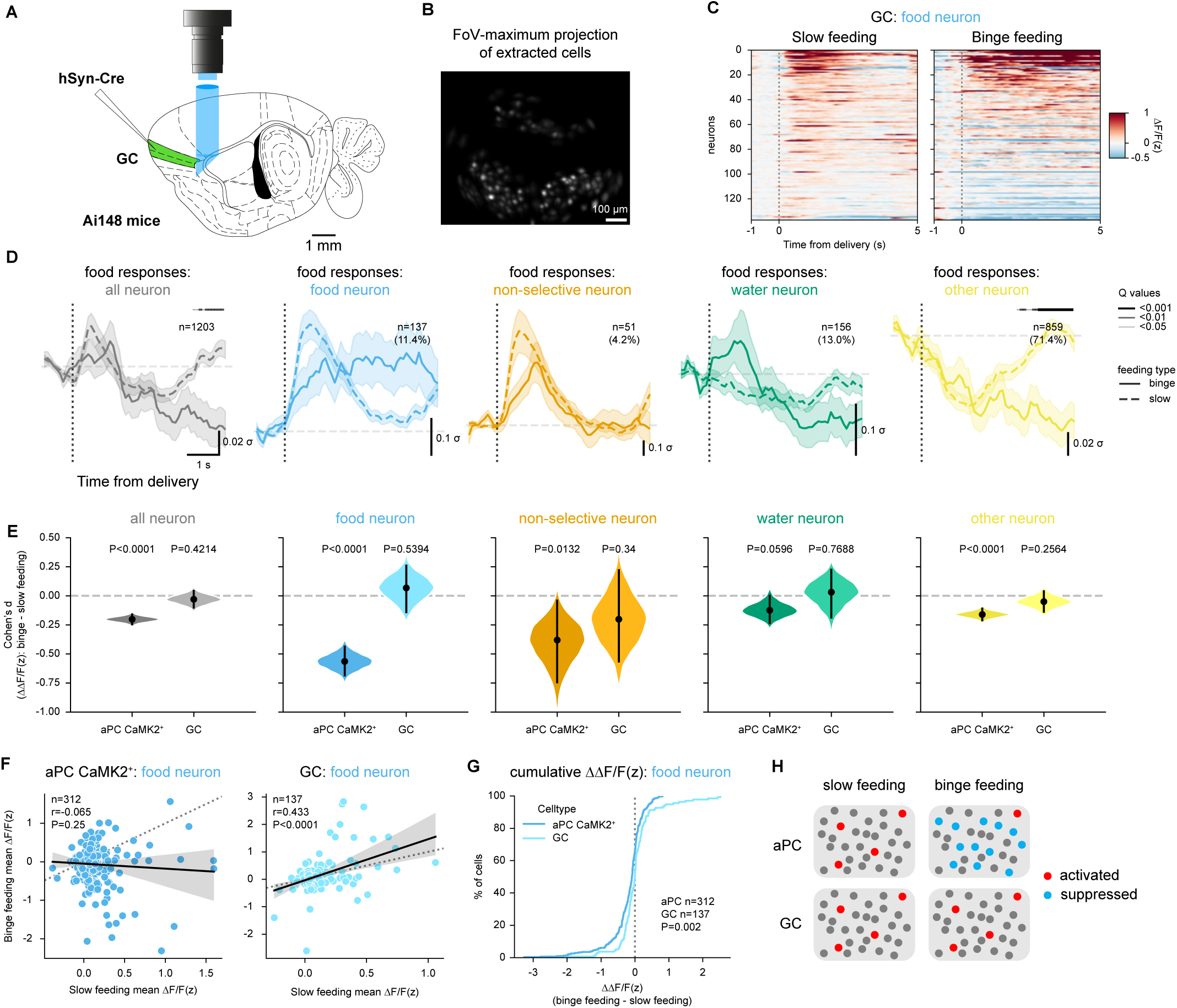
Flavor representation in the GC is stable across feeding rates. (**A**) Schematics of GRIN lens/Prism implantation in the GC. (**B**) Cell maps extracted by *CNMFe*. (**C**) Trial-averaged responses of individual GC food-activated neurons upon slow feeding and binge feeding (n=137 from 3 mice). (**D**) Trial-averaged responses of the whole population and subclasses of GC neurons upon slow and binge feeding (n=1203 cells from 3 mice). The shaded line above denotes the adjusted P-values (Q-values) of each time point, with different line widths representing different values (from thin to thick: Q <0.05, <0.01, <0.001). (**E**) Estimated effect size (Cohen’s d) of binge feeding-induced modulation in the aPC CaMK2^+^ and GC neurons within individual subclasses. P-values are calculated with the permutation test with 5000 times bootstrapping (n is the same as in **D**). (**F**) Cell-wise comparison of neuronal responses upon slow feeding and binge feeding in food-activated aPC CaMK2^+^ and GC neurons (n is the same as in Fig. 1F and in **C**). (**G**) Cumulative distribution of the difference (binge feeding vs. slow feeding) of z-scored ΔF/F (ΔΔF/F(z)) in food-activated aPC CaMK2^+^ and GC neurons (n is the same as in **F**). (**H**) Schematics of neuronal responses in the aPC and the GC during slow feeding and binge feeding. For (**D**), data are shown as mean ± s.e.m. For (**E**), data are shown as means of bootstrapped effect sizes (Cohen’s d) ± 95% confidence interval. For (**F**), r and P represent the correlation coefficient and P-value of Pearson’s r. For (**G**), the P-value is calculated from the Kolmogorov-Smirnov test.

To further analyze these distinct modes of flavor representation across the two cortices, we correlated the single-cell activity between binge and slow feeding in food-activated neurons in the aPC and the GC. In the GC, the activation level of food-activated neurons during slow feeding linearly correlates with the activity changes during binge feeding (Fig. 2F). In the aPC, however, we found no correlation between responses during slow and binge feeding, suggesting that binge-induced activity modulation is independent of the neuronal responses during slow feeding (Fig. 2F). The cumulative distribution of the amplitude difference between binge and slow feeding shows that in the GC, the net reduction is smaller and a larger fraction of cells show an increased binge feeding related neuronal response compared to the aPC (Fig. 2G). We interpret our finding as a non-uniform, general inhibition during binge feeding (Frank et al. 2019). Thus, flavor representation in the GC is preserved during binge feeding, whereas binge feeding induces a generalized activity suppression in the aPC (Fig. 2H).

### Binge feeding-induced anterior piriform cortex suppression is not inherited from the olfactory bulb

The aPC is the first cortical relay of the olfactory system and receives sensory afferent inputs from the olfactory bulb (Bekkers and Suzuki 2013). We tested if a reduction in sensory input from the olfactory bulb could explain global suppression during binge feeding in the aPC. To this end, we performed *in vivo* head-fixed 3-photon Ca^2+^ imaging in the olfactory bulb (OB) mitral cells during binge feeding. Mitral cells are the major neuron population propagating odor information to higher olfactory cortices, including aPC (Blazing and Franks 2020), and OB mitral cells remained activated upon binge feeding (Fig. 3A-C, S6A, B). At the population level, net excitatory OB output to the aPC was similar during slow and binge feeding (Fig. 3D). These findings align with the observation that intracortical connections and not the sensory afferents dominate neuronal activity in the aPC (Poo and Isaacson 2011). We conclude that binge feeding-induced aPC suppression is not inherited from the OB.

**Figure. 3.**
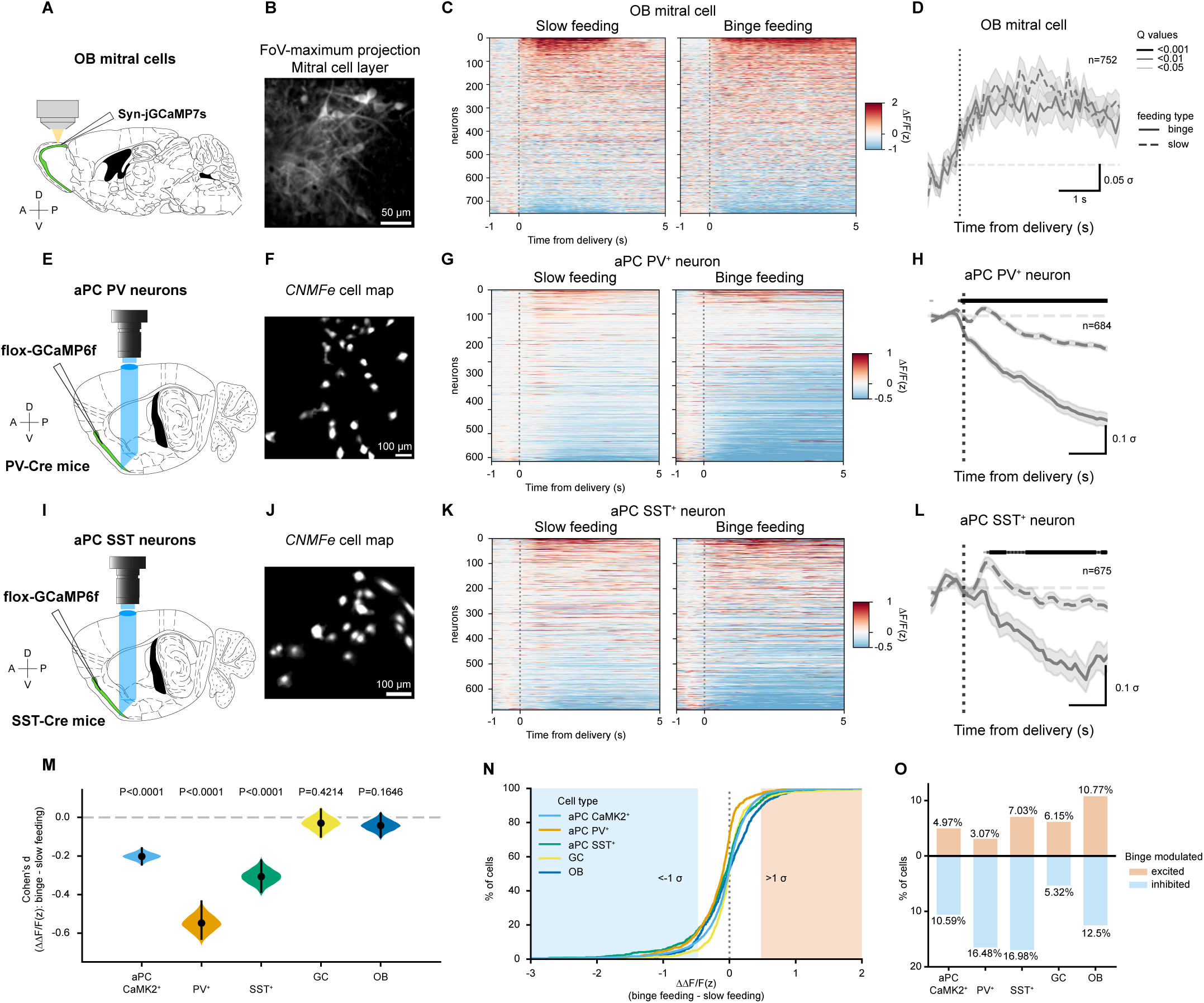
Binge feeding-induced aPC suppression is not inherited from the OB and extends to GABAergic aPC neurons. (**A**) Schematics of 3P-Ca^2+^ imaging in OB mitral cells. (**B**) FoV of OB mitral cells. (**C**) Trial-average of individual OB mitral cells upon slow and binge feeding (n=752 cells from 4 mice). (**D**) Trial-average of OB mitral cell population responses upon slow and binge feeding (n is the same as in **C**). (**E**) Schematics of Ca^2+^ imaging in aPC PV^+^ neurons. (**F**) Cell map of aPC PV^+^neurons extracted by *CNMFe*. (**G**) Trial-average of individual aPC PV^+^neurons upon slow and binge feeding (n=684 cells from 3 mice). (**H**) Trial-average of population aPC PV^+^neuron responses upon slow and binge feeding (n is the same as in **G**). (**I**) Schematics of Ca^2+^ imaging in aPC SST^+^neurons. (**J**) Cell map of aPC SST^+^ neurons extracted by *CNMFe*. (**K**) Trial-average of individual aPC SST^+^ neurons upon slow and binge feeding (n=675 cells from 3 mice). (**L**) Trial-averaged of population aPC SST^+^ neuron responses upon slow and binge feeding (n is the same as in **K**). (**M**) Estimated effect sizes of binge-induced modulation in aPC CaMK2^+^, aPC PV^+^, aPC SST^+^, GC, and OB mitral cells. (**N**) Cumulative distribution of binge-induced modulation of ΔF/F(z) in aPC CaMK2^+^, aPC PV^+^, aPC SST^+^, GC, and OB mitral cells. (**O**) Percentages of binge feeding modulated population in aPC CaMK2^+^, aPC PV^+^, aPC SST^+^, GC, and OB mitral cells. For (**D**), (**H**), and (**L**) data are shown as mean ± s.e.m, and the shaded line above denotes the adjusted P-values (Q-values) of each time point, with different line widths representing different values (from thin to thick: Q <0.05, <0.01, <0.001). For (**M**), data are shown as means of bootstrapped effect sizes (Cohen’ s d) ± 95% confidence interval.

### Binge feeding-induced anterior piriform cortex suppression extends to the major classes of local GABAergic neurons

Since the excitatory sensory drive is unaffected by feeding rate, enhanced recruitment of inhibitory interneurons during binge feeding could underlie the global suppression of the aPC. Local inhibitory feedback interneurons –activated by recurrent excitatory activity –predominantly inhibit odor responses in the aPC, whereas the contribution of feedforward interneurons is minor (Bolding and Franks 2018). We next probed the activity levels of aPC PV^+^ and SST^+^ inhibitory interneurons during slow and binge feeding with miniscope Ca^2+^ imaging, as they cover a large proportion of local feedback circuits (Large et al. 2016). Both PV^+^ and SST^+^ interneurons show strong suppression upon binge feeding. In contrast, the population responses of both types of interneurons showed net activity increases during slow feeding (Fig. 3E-L, S7A-F, S8A-C). Therefore, changes in the excitation-to-inhibition ratio of sensory afferent and local recurrent aPC circuits do not seem to mediate the global suppression of aPC activity during binge feeding. The binge feeding-induced suppression of activity in the aPC affects both the excitatory neurons and the major classes of local inhibitory interneurons. Therefore, binge feeding-induced suppression globally affects most local circuits in the aPC.

### Magnitude of binge feeding-induced anterior piriform cortex suppression correlates with appetite and depends on olfactory perception and metabolic state

Flavor perception of food items contributes to satiation and bypassing flavor perception via an intragastric catheter reduces satiation and accelerates gastric emptying of identical food items (Cecil, Francis, and Read 1999). The suppression of neuronal activity, specifically in the aPC described here, reduces the sensory representation of food items. Accordingly, we hypothe-sized that the aPC suppression during binge feeding could constitute a mechanistic link between a sensory neuronal response pattern in the flavor system and decreased satiation. Under *ad libitum* feeding conditions, mice consumed different amounts of food on different experimental days, which we take as a proxy for differences in satiation. This noticeable behavioral variability in our recording sessions correlates with temporal progression, suggesting binge feeding gradually escalates over time (Fig. S9A). We, therefore, investigated whether this behavioral variability maps onto the aPC neuronal responses. Using a linear mixed model, we found a robust time-independent correlation between the initial binge eating-induced aPC suppression and subsequent food consumption on each recording session (Fig. 4B). Suppression was always quantified during the onset of a binge bout within the first 4 seconds after initiation. Our model, therefore, quantifies suppression independently of feeding duration. The strong correlation between neuronal activity patterns and food intake does not exist for slow feeding aPC responses (Fig. 4B, C). We also observed consumption-correlated suppression of neuronal activity in aPC PV^+^ neurons. In contrast, aPC SST^+^ neurons and GC neurons did not show a correlation between neuronal activity and food intake. (Fig. S9C). These findings suggest that consumption-correlated neuronal modulations are mostly restricted to the olfaction component of flavor perception.

**Figure. 4.**
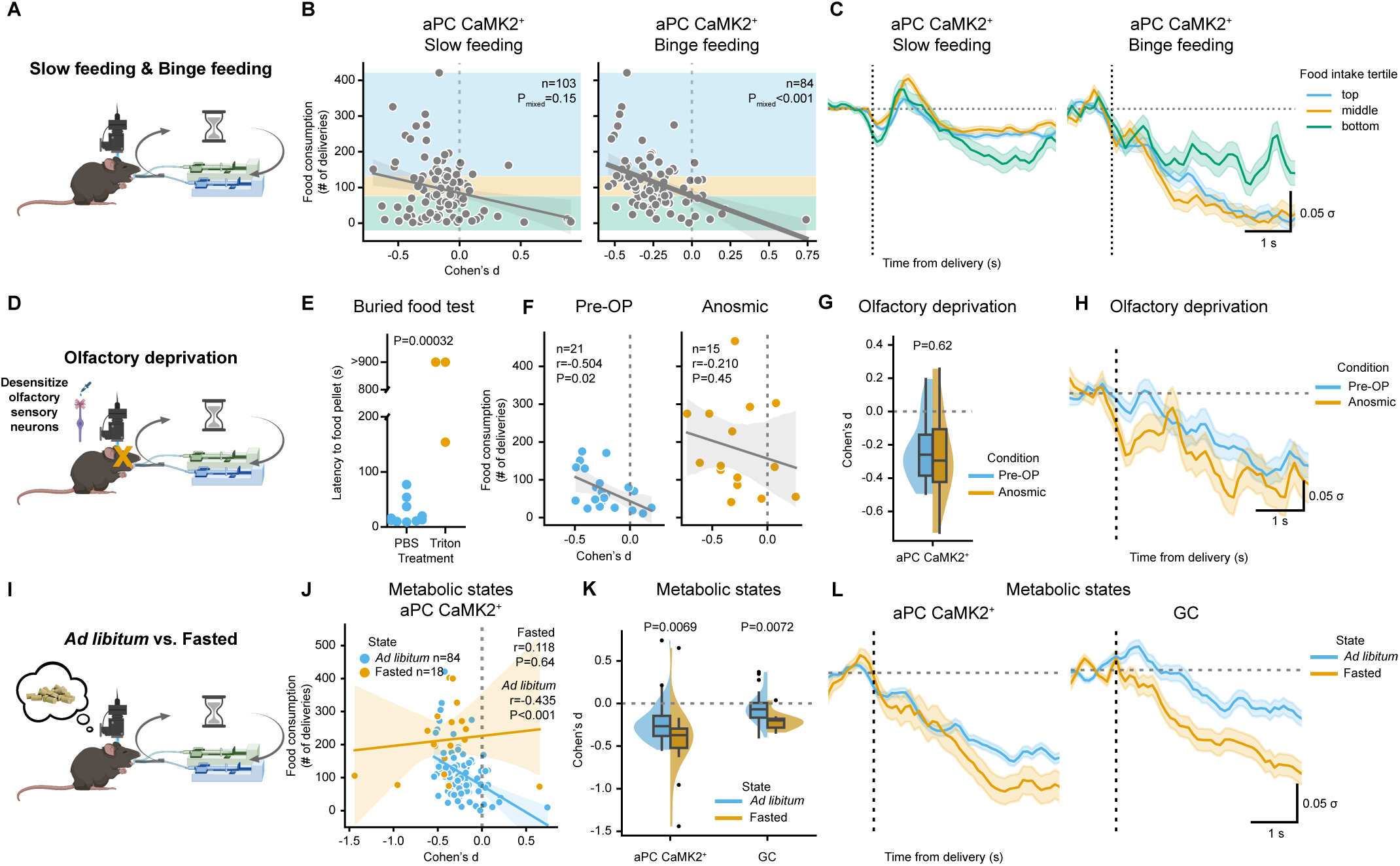
Magnitude of binge feeding-induced anterior piriform cortex suppression correlates with appetite and depends on olfactory perception and metabolic state. (**A**) Schematics of feeding and Ca^2+^ imaging paradigm. (**B**) Correlations between session-specific food intake and modulation of neuronal activity in aPC CaMK2^+^ neurons upon slow and binge feeding (n=103 slow feeding sessions and 84 binge feeding sessions from 8 mice). (**C**) Trial-average responses of aPC CaMK2^+^ neurons upon slow and binge feeding clustered by food intake (n=34, 35, 34 sessions for top, middle, bottom clusters in slow feeding from 8 mice, n=28, 28, 28 sessions for top, middle, bottom clusters in binge feeding from 8 mice). (**D**) Schematics of anosmic paradigm. (**E**) Latency of mice finding the buried food pellet 48 hrs after treatment (n=10 mice for PBS treated group, n=3 for Triton treated group). (**F**) Correlations between session-specific food intake and modulation of neuronal activity in aPC CaMK2^+^ neurons upon binge feeding with intact olfaction and under anosmic conditions (n=21 Pre-OP sessions and n=15 Anosmic sessions from the same 3 mice). (**G**) Binge-induced modulation of neuronal activity with intact olfaction and under anosmic conditions (n is the same as in **F**). (**H**) Trial-averaged activity of aPC CaMK2^+^ neurons upon binge feeding with intact olfaction and under anosmic conditions (n is the same as in **F**). (**I**) Schematics of fasting paradigm. (**J**) Correlations between session-specific food intake and modulation of neuronal activity in aPC CaMK2^+^ neurons upon binge feeding under *ad libitum* or overnight fasted conditions (n=18 fasted sessions from 5 mice, n for *ad libitum* conditions is the same as in **B**). (**K**) Binge-induced modulation of neuronal activity under *ad libitum* or overnight fasted conditions (n is the same as in **J**). (**L**) Trial-averaged responses of aPC CaMK2^+^ neurons upon binge feeding under *ad libitum* or overnight fasted conditions (n is the same as in **J**). For (**C**), (**H**), and (**L**) data are shown as mean ± s.e.m. For (**F**) and (**J**) r and P represent the correlation coefficient and P-value of Pearson’s r. For (**B**) P represents the significance level of Cohen’s d to food consumption by a linear mixed model. For the box plot in **G** and (**K**) the center line shows the median, the box limits show the quartiles, the whiskers show 1.5x the interquartile range, and the points show the outliers.

We further examined the necessary factors for the consumption-correlated generalized suppression of aPC neuronal activity during binge feeding. To test whether the olfactory perception of the food items is a necessary for consumption-correlated suppression of neuronal activity, we performed nasal lavage with 0.5% Triton solution to induce temporary anosmia in mice (Cummings et al. 2000) (Fig. 4D, E, anosmia verified by buried food test). We performed imaging experiments before and after the intervention. Our procedure caused a 30% drop on average in the number of active neurons in the field of view detected with imaging, consistent with reduced olfactory inputs to the aPC (Fig. S9B). We then compared the correlation of the aPC activity suppression to consumption in the same mice before and after the anosmia-inducing treatment. In contrast to the pre-anosmia condition, anosmia-inducing treatment abolishes the correlation between the suppression of aPC neuronal activity and consumption (Fig. 4 F, G). Under anosmia, we still observed binge eating-induced aPC suppression (Fig. 4H) in the presence of generally enhanced food intake (Fig. 4F). This result suggests that intact sensory olfactory perception is a prerequisite for binge eating-induced aPC suppression correlated to consumption. Metabolic states like hunger and satiety profoundly affect sensory systems (Soria-Gómez et al. 2014; Aimé et al. 2007; Freeman 1960; Prud’homme et al. 2009; Albrecht et al. 2009). Consequently, we wondered if changes in the metabolic state affect the consumption-correlated suppression of aPC neuronal activity during binge feeding. We altered the metabolic state of mice by overnight fasting. We found enhanced food intake in these mice and enhanced binge feeding-induced suppression of aPC neuronal activity compared to *ad libitum*-fed conditions (Fig. 4I-L). Under fasting, the correlation between food consumption and aPC neuronal activity suppression is lost (Fig. 4J). An increase in suppression was also observed in the GC of fasted mice (Fig. 4K, L). We conclude that the suppression of neuronal activity during binge feeding in the aPC is enhanced by fasting and is also observed in the GC.

### Optogenetically suppressing anterior piriform cortex neurons promotes feeding

We have so far established the correlative relationship between feeding rate, olfactory flavor representation, and metabolic state. While binge feeding and appetite clearly covary with global activity suppression in the aPC, it is unclear if the sensory effect we observe is an epiphenomenon (***H0***, Fig. 5A) or reciprocally interacts with feeding behavior in a feedback loop (***H1***, Fig. 5A). We, therefore, next asked whether there was a causal relationship between suppressed aPC activity and feeding behavior by inhibiting aPC during feeding. This tested if aPC suppression alone is sufficient to increase food consumption (***H1***, Fig. 5A). We employed a closed-loop optogenetic inhibition paradigm to silence aPC excitatory neurons at the initiation of binge feeding bouts (Fig. 5B, E). To suppress activity at the behavioral timescale (tens of seconds) of binge feeding bouts while minimizing the illumination period, we chose the highly light-sensitive mosquito opsin eOPN3 (Mahn et al. 2021) to provide long-lasting suppression of recurrent excitatory fibers in the aPC (Fig. S10C for optical fiber implant coordination). We found that mice consumed more food when aPC activity was optogenetically suppressed upon feeding (Fig. 5F-I). The optogenetic suppression of aPC activity prolonged the individual feeding bouts, while the number of feeding bouts remained similar, suggesting that optogenetic aPC suppression predominantly affects consummatory and not appetitive behavior (Fig. 5F and Fig. S10A, B). Light stimulation alone did not affect feeding behaviors in control mice transduced with AAVs encoding tdTomato (Fig. 5G-I). Our data infers that binge feeding-induced aPC suppression is causally linked to feeding behaviors, suggesting a functional role of binge feeding-induced aPC suppression in modulating appetite (***H1***, Fig. 5A)

**Figure. 5.**
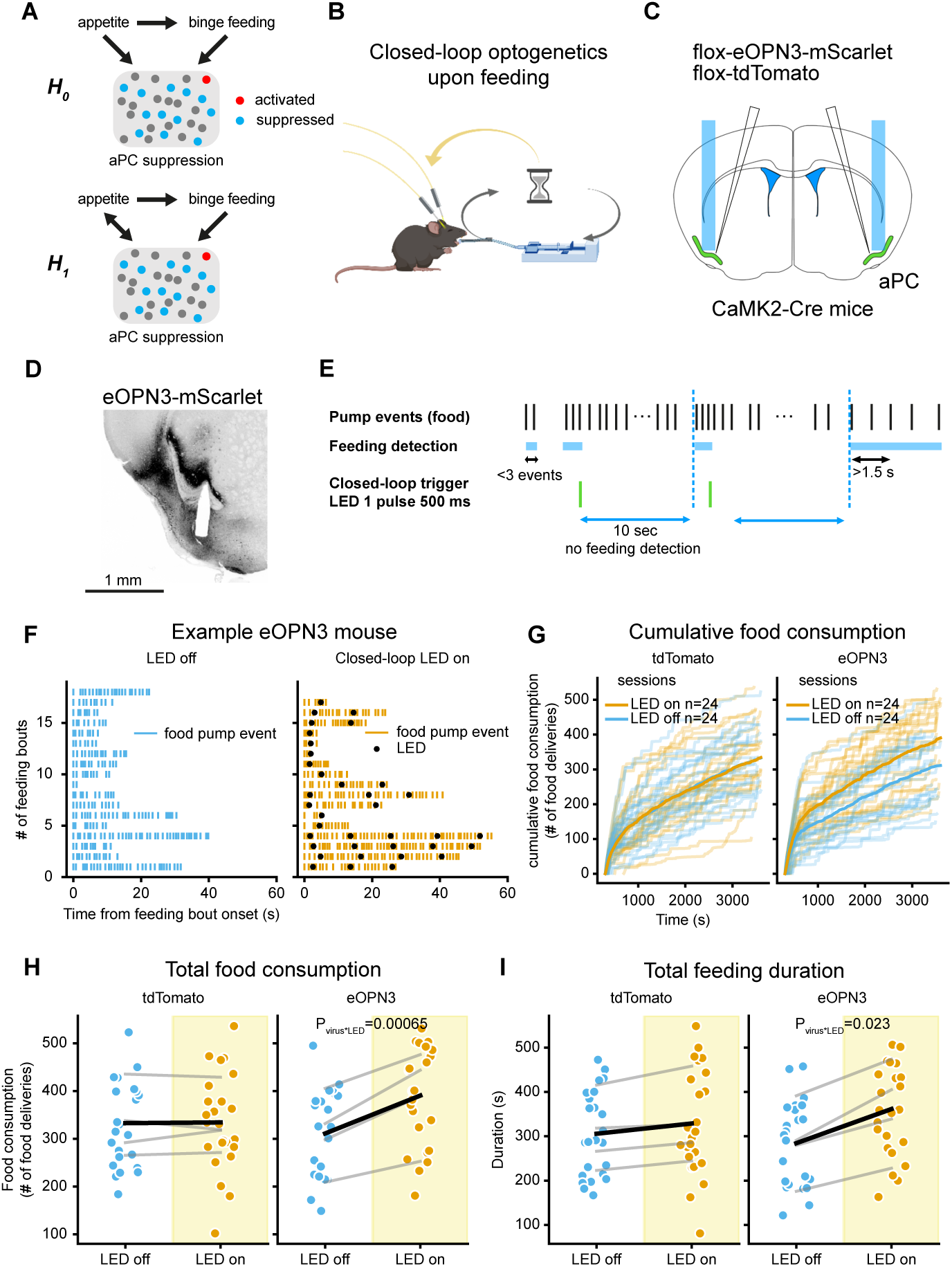
Optogenetically suppressing aPC neurons promotes feeding. (**A**) Schematics of hypotheses on relationships of appetite, binge feeding and aPC suppression. (**B**) Schematics of closed-loop optogenetics experiment setup and feeding paradigm. (**C**) Schematics of viral injection and optical fiber implants bilaterally in the aPC. (**D**) Brain slice with optical fiber implant path. (**E**) Schematics of feeding-based closed-loop optogenetics paradigm. (**F**) Example feeding bouts in an eOPN3-expressing mouse without light stimulation (left panel) and with closed-loop light stimulation (right panel). (**G**) Cumulative feeding events through experimental sessions for mice expressing tdTomato (left panel, n=12 LED off and 12 LED on sessions from 4 mice) and mice expressing eOPN3 (right panel, n=12 LED off and 12 LED on sessions from 4 mice). (**H**) Effects of light stimulation on total food consumption in tdTomato- and eOPN3-expressing mice (n is the same as in **G**). (**I**) Effects of light stimulation on total feeding duration in tdTomato- and eOPN3-expressing mice (n is the same as in **G**). For (**H**) (**I**) grey lines denote the data from the same mice, and the black line denotes the mean. The P-values are calculated from a linear mixed model (see **Methods**).

## DISCUSSION

The role of olfactory flavor representation during feeding behaviors is poorly understood despite its essential contribution to flavor experience (Shepherd 2006; Maier et al. 2015). Using cell-type specific *in vivo* Ca^2+^ imaging and optogenetics in freely behaving mice, we provide circuit-level evidence that suppression of flavor representation in the aPC during binge feeding actively enhances food intake. While chronic effects of olfactory alterations on food intake and metabolism have been reported (Riera et al. 2017; Tucker, Overton, and Fadool 2012), we found an acute functional role of olfaction at the level of individual feeding bouts. Our findings suggest that the olfactory representation of flavor during feeding provides feedback for sensory satiation to modulate food intake and homeostasis in real-time.

Metabolic states have profound effects on sensory systems, especially olfaction. Hunger increases neuronal and behavioral responses to odors, and in contrast, satiety decreases them (Soria-Gómez et al. 2014; Aimé et al. 2007; Freeman 1960; Prud’homme et al. 2009; Albrecht et al. 2009). Sensory detection of food items leads to a rapid switch in activity patterns from consumption-promoting AgRP neurons to consumption-inhibiting POMC neurons in the arcuate nucleus of the hypothalamus. The amplitudes of these foraging-related switches in hypothalamic activity are enhanced by fasting and sensory signals from food items with a high hedonic value (Chen et al. 2015; Betley et al. 2015; Mandelblat-Cerf et al. 2015). We demonstrated that the amplitude of binge feeding-induced suppression in the aPC reflects internal appetite levels and is enhanced by fasting, supporting that metabolic states strongly modulate olfactory representation during feeding.

The sensory experience of food items contributes to satiety. Bypassing sensory experiences of food within the oral cavity by direct gastric infusion reduces satiety and accelerates gastric emptying compared to regular feeding in humans and rodents (Berkun, Kessen, and Miller 1952; Cecil, Francis, and Read 1999; Stratton and Elia 1999). A recent study also demonstrates a brainstem circuit of gustatory oral sensory feedback that induces satiation (Ly et al. 2023). Our study supports the idea that suppressed olfactory flavor representation, via direct binge feeding or artificially suppressing the aPC, constitutes an acute intrinsic behavior-associated cortical mechanism leading to more consumption, inferring lower satiety levels. The reciprocity between feeding behavior and olfactory representation demonstrated here (Fig. 5a) extends the emerging theory that perception and action reciprocally interact (Buzsáki 2019).

From an evolutionary perspective of food scarcity, overeating induced by suppression of sensory satiety would be pro-survival. In the surplus of food environment human beings are facing today, an increased eating rate is commonly linked to overeating (Hall et al. 2019) and obesity (Ohkuma et al. 2015), while reducing the eating rate can effectively mitigate food consumption (Hurst and Fukuda 2018; Scisco et al. 2011; Bolhuis et al. 2014). Here, we provide evidence that feeding rate-dependent modulation of olfactory flavor representation modulates appetite (Fig. S11). Our findings add the perspective of sensory experience and awareness to the notion that it is not only what you eat but also how you eat.

## ACKNOWLEDGEMENTS

We thank Susanne Rieckmann, Monika Dopatka, Anny Kretschmer, Katja Czieselsky and Celina Ernst for their excellent technical assistance; Daniel Parthier for his inputs on statistical analysis; Yen-Chung Chen for his inputs on programming; Melissa Long from the Charité Animal Behavioral Phenotyping Facility, Charité Advanced Medical BIOimaging Core Facility, and Charité Viral Core Facility for their assistance; James Poulet and David Owald for their feedback on the manuscript; Robin Blazing and Kevin Franks for sharing *in vivo* optical fiber implantation protocols; Andreas Klaus for his input at the early stage of this study.

HL is supported by the PhD Fellowship from the Einstein Center for Neurosciences Berlin and the Taiwanese Government Scholarship to Study Abroad. FWJ is supported by the Deutsche Forschungsgemeinschaft (DFG) (project 458236353 and project 520223756 - SPP2411 LOOPs). BRR is supported by the DFG (project 327654276 - SFB1315 and project 273915538 - SPP1926). DS is supported by the DFG (project 184695641 –SFB 958, project 327654276 –SFB 1315, project 415914819 –FOR 3004, project 431572356, the German Excellence Strategy EXC-2049-390688087, NeuroCure), by the European Research Council (ERC) under the European Union’s Horizon 2020 research and innovation program (BrainPlay Grant agreement No. 810580) and by the Federal Ministry of Education and Research (BMBF, SmartAge –project 01GQ1420B).

## AUTHOR CONTRIBUTIONS

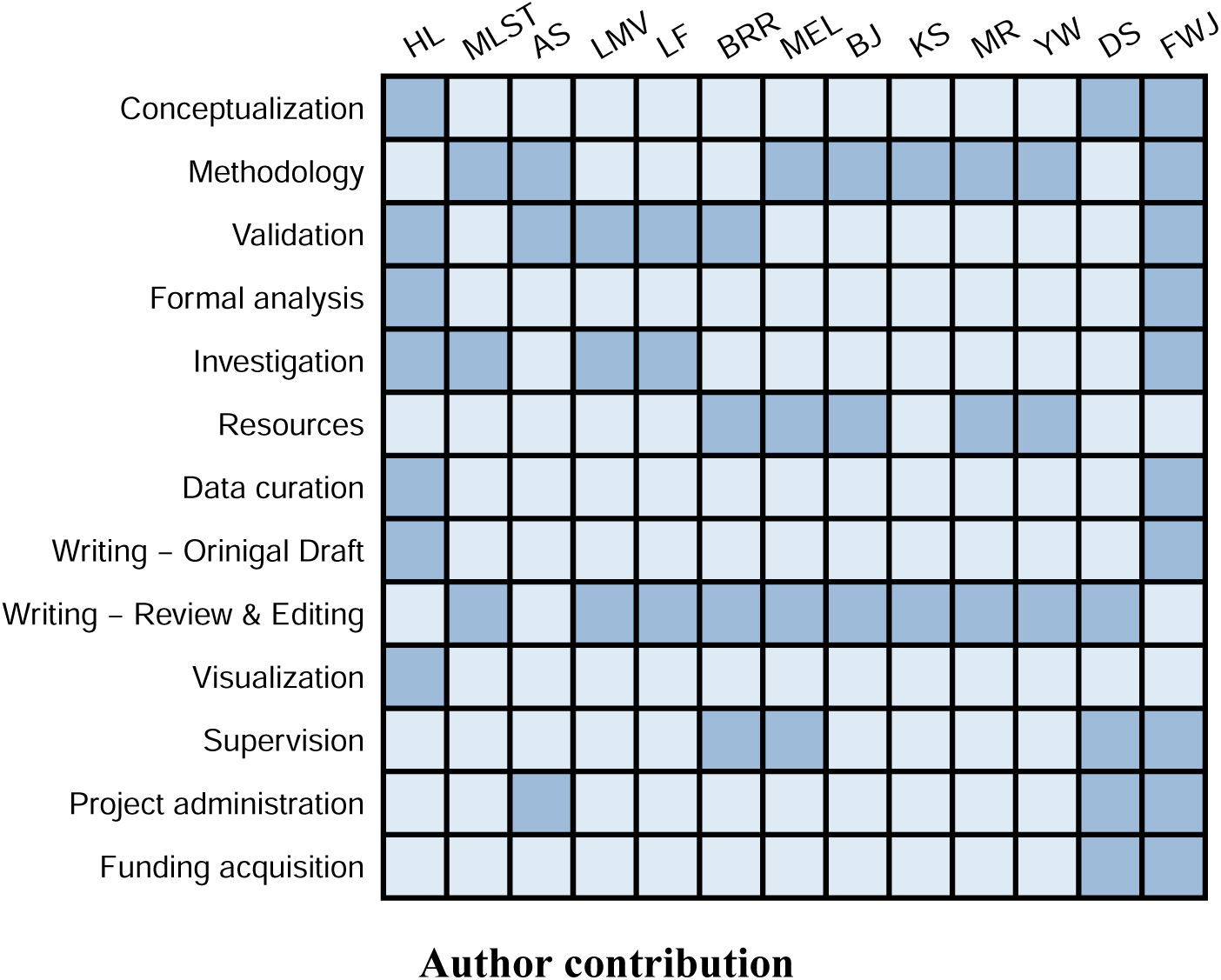

Conceptualization: HL, DS, and FWJ; methodology: AS, FWJ, KS, MR, YW, MLST, BJ, and MEL; validation: AS, HL, LF, LMV, BRR, and FWJ: formal analysis: HL and FWJ; investigation: HL, FWJ, MLST, LF, and LMV; resources: MEL, BJ, MR, YW, DS, and BRR; data curation: HL and FWJ; writing - original draft: HL and FWJ; writing - review & editing: DS, BRR, MR, BJ, KS, MLST, MEL, LF, LMV, and YW; visualization: HL; supervision: FWJ, DS, MEL, and BRR; project administration: AS, FWJ, and DS; funding acquisition: FWJ and DS.

## DECLARATION OF INTERESTS

YW owns PhenoSys equity. Other authors have declared no competing interest.

## DATA AND CODE AVAILABILITY

Source data for individual figure panel can be found on a GitHub repository (https://github.com/hung-lo/BingeFeeding_2023). Raw data will be available upon request due to the large file size (∼2-3 TB). Code for plotting individual figure panel can be found on a GitHub repository (https://github.com/hung-lo/BingeFeeding_2023). All code for data processing and analysis will be deposited to a GitHub repository upon acceptance of this manuscript.

## METHODS

### Animals

Animals were kept at the animal facility of Charité, under a regular 12/12 hour light-dark cycle. All procedures involving animal experiments were approved by the local authorities and ethics committee (LaGeSo Berlin, license numbers G0278/16, G0313/16, and G0156/20). To image excitatory neurons in the aPC, we cross-bred Ai93D mice, Rosa-tTA mice, and CaMK2-Cre mice to obtain Ai93D; Rosa-tTA; CaMK2-Cre mice that express GCaMP6f in excitatory cells. To prevent early expression of GCaMP during development, the breeding pairs and offspring were fed with Doxycycline-containing food to suppress the expression of GCaMP6f until weaning. Due to suboptimal GCaMP6f expression in the GC in the abovementioned transgenic mice, we injected the AAV virus carrying hSyn-Cre in the GC of Ai148D mice to express GCaMP6f in the GC. To image OB mitral cells, we inject Syn-jGCaMP7s AAV virus in C57BL/6 mice. To image GABAergic neurons in the aPC, we performed viral injection of Cre-dependent GCaMP6f in PV-Cre or SST-Cre mice. For optogenetic experiments, CaMK2-Cre mice were bilaterally injected with Cre-dependent eOPN3 virus or a Cre-dependent tdTomato expressing virus for controls. All experiments including Ca^2+^ imaging and optogenetics are performed between 9 a.m. to 6 p.m. under regular light.

### Liquid food delivery system

To reduce stress, we performed experiments inside the animals’home cages. Cages were modified so that we could protrude the motorized lick spout (PhenoSys, Berlin, Germany) into the cage. After a 5-minute baseline period, motorized lick spouts were presented in the cage and primed for liquid delivery for 2 min with 1 min intervals between presentations. During these intervals, the spouts were retracted. For olfactory isolation, we presented the lick spout inside a glass tube with an opening for animals to reach the spout. Air suction from the glass tube limited olfactory responses to the food odor to the time period just before and while mice interacted with the lick spout. Lick spouts were equipped with piezo sensors to register each licking. Licks triggered a 400-ms activation of electrical pumps, which resulted in the delivery of one droplet (∼1.8 µL) of strawberry or chocolate-flavored Ensure (Abbott Laboratories) or water.

After delivery, we set a refractory delay period of 4 sec (slow feeding) or 400 ms (binge feeding). During slow feeding, we provided Ensure and water at a ratio of 7 to 3. Four slow-feeding rounds and three binge-feeding rounds were interchanged in a pseudorandom order. Mice were *ad libitum* fed before the experiments, with a maximal period of up to 4 hours of pre-experiment food deprivation during the light cycle. Fasting was performed once a week for 20-22 hours before starting the experiment.

### Surgery procedures

#### Stereotactic injection

Mice were anesthetized by inhalation anesthesia with isoflurane (induction: 4-5%, then 1-2% with oxygen, flow rate 0.5-1 l/min). Mice were local anesthetized with Lidocaine (1-2%) subcutaneous injection preincision. A craniotomy was performed over the stereotactically determined target regions (Table. S1) using a semi-automatic neurostar stereotactic apparatus (Neurostar, Tübingen, Germany). The virus (0.4 to 1 µL) was injected using a 10 µL-Hamilton syringe. Postoperative pain was prevented by Carprofen (5 mg/kg) subcutaneous injection right before surgery and in the first 3 days after surgery. After the surgery, the animals recovered for at least two weeks. In some experiments, implantation of the prism or the optic stimulation fiber was performed right after viral injection.

#### GRIN lens implantation

Mice (>P50) were anesthetized for the procedure by inhalation anesthesia with isoflurane (induction: 4-5%, then 1-2% with oxygen, flow rate 0.5-1 L/min). Mice were local anesthetized with Lidocaine (1-2%) subcutaneous injection preincision. The anesthetized animals were fixed in the stereotact (Neurostar, Tübingen), and a craniotomy was performed over the stereotactically determined target region (Table. S1). The side length of the quadratic craniotomy was slightly larger than the side length of the prism base, approximately 1.2 mm.The insertion tract was paved by aspiration of brain tissue until ∼1 mm above the image plane of the microscope. Aspiration was performed through a thin needle (23G, sharp end) linked to a vacuum pump, the procedure was performed twice to ensure sufficient aspiration of brain tissues. Any small hemorrhagic foci that occurred were staunched by Gelfoam. After removal of the Gelfoam and any pending blood clots, insertion of the microendoscopic lens (GRIN lens attached with a Prism (1 mm diameter, ∼9.1 or ∼4.3 mm long, Inscopix) to the desired image plane is performed at a rate of 100 µm/min according to the coordinates in Table S1. With the aid of an adhesive (VetBond, 3M or TRUGLUE, TRUSETAL) and dental cement (Super-Bond C&B, SUNMEDICAL), microendoscopes with attached baseplates (Inscopix) were fixed to the skull, and the optical surface was protected from contamination by a plastic cap (Inscopix baseplate cover). Postoperative pain was prevented by Carprofen (5 mg/kg) subcutaneous injection right before surgery and in the first 3 days after surgery. In a subset of mice, we did not use microendoscopes with attached baseplates. In this case, the slightly protruding microendoscope was fixed to the skull with adhesive and dental cement, and the optical surface was protected from contamination using a silicone cap. In these mice, a baseplate was fixed in the desired optical plane above the protruding lens in a surgery that was performed at a minimum of four weeks after the lens implant.

#### Cranial window for olfactory bulb imaging

Mice (C57BL/6, <P40) were anesthetized with isoflurane (induction: 4-5%, then 1-2% with oxygen, flow rate 0.5-1 l/min). Mice were local anesthetized with Lidocaine (1-2%) subcutaneous injection preincision. After the scalp and periosteum were removed, a 3mm craniotomy was made over the two bulb hemispheres. An injection micropipette (tip diameter, 10–20 *µ*m) was filled with AAV1.Syn.jGCaMP7s.WPRE virus solution (Penn Vector Core), and 100 nl was injected 50 nl/min at a depth of 300 *µ*m in either bulb hemisphere (see Table S1 for coordinates). After injection, a semi-circular <3 mm stack of two glass coverslips, glued to each other using optical adhesive, was fitted into the craniotomy and sealed with cyanoacrylate glue and dental cement. Finally, a light-weight head-post was fixed on the skull over the left hemisphere with light-curing adhesives (RelyX, 3M) and dental cement (Ortho-Jet, Lang Dental). Postoperative pain was prevented by Buprenorphine (0.05-0.1 mg/kg) and Carprofen (5 mg/kg) subcutaneous injection right before surgery and then Carprofen (5 mg/kg) in the first days after surgery. Head-fixed 3-photon imaging experiments began 3 weeks after the virus injection.

### Ca^2+^ imaging

#### Habituation

For freely-moving recordings, before starting the combined behavioral and imaging sessions using Ensure and water, mice were habituated to the lick spout delivery with 10% sucrose solution. Mice had to reach a criterion of 25 sucrose deliveries in slow feeding mode in a 45-minute habituation session before the actual measurements began. Mice were further habituated with additional air suction around the lick spout and dummy scope mounting once they had learned to drink from the lick spout. The habituation period usually lasted ∼3 weeks.

For head-fixed recordings, the habituation of mice to head-fixation began at least 5 days prior to imaging. On the first day, the animal was head-fixed on a running wheel for 5 min and then gradually increased each day until it was calm for 1 hour. At least one day before imaging, a lick spout with milk/water within easy reach for licking was introduced.

#### in vivo imaging: miniscope

The miniaturized microscope (nVista miniscope, Inscopix, CA, USA) was mounted right before the imaging session started without anesthesia. Before the recording started, mice were allowed to explore the home cage for 3-5 minutes. After a baseline period of 5 min, the lick spout protruded according to the protocol described above. For each mouse, the imaging settings (LED intensity, gain, focus⋯etc.) were individually tuned to reach a similar level of brightness (mean values around 50-60 A.U. in fluorescence histogram function in Inscopix acquisition software). We recorded at 20 Hz with a single focal plane. Most imaging sessions were 4x spatially down-sampled during acquisition to save storage space. The behavioral system was linked with the Inscopix system using TTL pulses upon pump activation. We performed up to 25 imaging sessions per mouse across 5 weeks, and overnight food deprivation was performed once per week.

#### In vivo imaging: 3-Photon

Imaging from head-fixed mice was performed with a home-built 3-photon microscope. The laser (Opera-F, pumped by Monaco, Coherent) provided light pulses at 1300 nm wavelength and 1 MHz repetition rate for excitation of jGCaMP7s. The laser output passed a four-pass prism pulse compressor for dispersion compensation. Laser power was adjusted using a motorized half-wave plate and a polarizing beam splitter, and was below 20 mW under the objective. We used a Nikon25x/1.1 objective and dual linear galvanometers at a frame rate of ∼10Hz. Image acquisition was synchronized with laser pulses and was controlled by LSMAQ. Time-series images (200 x 200 pixels) were recorded at depths of 200-300 *µ*m below pia at the mitral cell layer.

### Triton X-100 application

Nasal lavage with 0.5% of Triton X-100 (in 0.1M PBS) can introduce temporal anosmia in mice for up to 3 weeks (Cummings et al. 2000). Mice were anesthetized with Ketamin (100 mg/Kg), Xylazine (20 mg/Kg) and Acepromazine (3 mg/Kg) intraperitoneal injection. They received Caprofen (5mg/ Kg) subcutaneous injection before and the day after the Triton X-100 (experimental group) or PBS (control group) applications. We applied 40 µL 0.5% Triton X-100 solution to each nostril with a gel loading pipette tip that was advanced for 2-3mm into the nostril. Triton solution was slowly applied with a micropump (Narishige, Japan) over several minutes on each side, with an interval of 5 min between the two nostrils. Foam building up at the opening of the nostrils was an indicator of a successful procedure. Throughout the procedure and until waking up from anesthesia, mice were kept on an inclined plane so that their nostrils were below their trachea and lungs in a heated chamber.

### Buried food test

To test the efficacy of Triton-induced anosmia, we examined mice’s ability to find a hidden food pellet located 1 cm deep in the bedding of the experimental cage. Mice were overnight food-deprived and habituated to the experimental cage for at least 5 minutes before the experiment started. After the food pellet was buried, mice were transferred into the experimental cage. The time to find the pellet was documented by the experimenter. If the mice did not find the pellet after 15 minutes, the experiment was stopped. Control experiments were performed with mice that had undergone the same lavage procedure with 0.1M PBS nasal lavage. The buried food tests were repeated every week to monitor mice’s smell ability to ensure mice remained anosmic throughout our experiments.

### Optogenetic experiments

Mice expressing eOPN3 in aPC excitatory neurons were anesthetized for the procedure by inhalation anesthesia with isoflurane (induction: 4-5%, then 1-2% with oxygen, flow rate 0.5-1 L/min). The anesthetized animals were fixed in the stereotact (Neurostar, Tübingen), and a craniotomy was performed over the stereotactically determined target region. Fiberoptic cannulas (200 µm diameter, NA 0.66, length 5mm, Doric lenses) were inserted bilaterally until reaching the target coordinates (Fig. S10c). The slightly protruding fiber with an attached zirconia sleeve for taking up the stimulation fiber patchcord was fixed to the skull with adhesive and dental cement, and the optical surface was protected from contamination using a plastic cap. Before starting the combined behavioral and optogenetic stimulation sessions using Ensure, the mice were habituated to the lick spout delivery with 10% sucrose solution. Mice had to reach a criterion of 25 sucrose deliveries in binge feeding mode and needed to successfully trigger sham closed-loop stimulations in a 20-minute habituation session before the actual measurements began. At the start of the experiment, we plugged a splitter branching patchcord (200 µm diameter, NA 0.57, Doric) connected to a mono fiberoptic patchcord (480 µm diameter, NA 0.63, Doric) onto the fiberoptic cannulas. Light from a Ce:YAG fiber light source (Doric) was delivered at an intensity of 8mW at the tip of the fibers. For these experiments, mice were granted constant access to the lick spout in the binge feeding mode over a period of 60 min. Upon detection of a binge bout (3 pump deliveries with a maximum of a 1.5 s interdelivery interval), a 500 ms light stimulus was delivered on “LED on”days. After the detection of a binge bout with a subsequent light stimulus, there was a minimal refractory period of 10 seconds until a binge bout could initiate the next light stimulus. Everything was similar on “LED off”days, apart from not stimulating with light upon binge bout detection. LED on and off days were alternated for 12 subsequent days.

### Imaging processing

Ca^2+^ movies obtained from miniscope recordings were first temporally downsampled to 10 Hz. We then cropped out regions in the field of view (FoV) where no active Ca^2+^ transients were visible. The same FoV cropping parameters were used throughout recordings from the same mice. Movies were then bandpassed with a spatial filter (low cutoff=0.005, high cutoff=0.500, Inscopix IDPS) and motion corrected (aligned to mean image or first frame, max_translation=20, Inscopix IDPS). Ca^2+^ traces were extracted with an adapted version of *CNMFe* (Zhou et al. 2018) from Inscopix (see Table S4) with the following parameters (Cell diameter: 10 px, PNR: 10 for excitatory cells and 20 for GABAergic cells, Corr: 0.8). Ca^2+^ traces were manually curated with predefined selection criteria (peak amplitude >80 A.U., baseline drifts less than 20% of peak fluorescence, clear cell shape, locate outside blood vessels, minimal motion artifacts of given regions of interest).

Image stacks from 3P imaging were loaded into Suite2P (Pachitariu et al. 2016) for motion correction, region-of-interest (ROI) segmentation and trace extraction using the settings specified in the appended exemplary .ops file. We used the ‘mean img’, ‘correlation map’, and ‘max projection’views of Suite2P to manually check and sort somatic from non-somatic ROIs of mitral cells. The output from Suite2P was analyzed in Python: Detected neuropil signals were subtracted. Remaining frames with movement artifacts were then detected and excluded based on the presence of post-registration x- and y-shifts at each time point. A further criterion was the phase correlation of individual frames and the reference image below a threshold of 50% of the maximum peak of phase correlation in the respective stack. After that, ΔF/F values and z-scores were calculated in customized Python scripts.

### Data analysis, statistics and plotting

All data analysis and statistics were performed using custom scripts in Python and R. Most figures were plotted in Python (matplotlib, seaborn) and figure and font sizes were later modified in Illustrator (Adobe). We used the Okabe-Ito color palette (Ichihara et al. 2008) to increase the accessibility of common forms of color blindness.

#### Data synchronization

Ca^2+^ imaging data and behavioral data were synchronized by finding the time lag of maximum cross-correlation between pump events in digital values from the PhenoSys behavioral protocol and the binarized pump-triggered TTL pulses recorded in the Inscopix system by Pynapple.cross_correlogram (Python).

#### Binge feeding bout detection and slow feeding processing

To detect binge feeding bouts, inter-pump intervals were calculated for each pump event, and only pump deliveries with intervals shorter than 2 s qualified as part of a feeding bout. Additionally, each feeding bout was required to include at least 3 pump deliveries.

Pump events in slow feeding mode were filtered out if no further lick event followed the initial lick event triggering pump activation. Initial motor artifacts from the movement of the lick spout were also removed from the further analysis. Since binge feeding bouts were guaranteed to have subsequent lick events by design, the exclusion of pump events following no lick events was not applied to binge feeding pump events.

#### Statistical analysis

Statistical analysis was performed in Python (Scipy, Numpy, Dabest) and R (Lmer). Individual statistical tests are listed under the respective figure legends and all statistical details are listed in the statistical summary Table S5. All statistical tests are performed with two-sided testing. Mean ± standard error of the mean (s.e.m.) or 95% confidence of interval were used to report statistics in figures. Applied statistical tests and the sample size for each analysis are listed in the figure legends and the statistical summary table. A significance level of p <0.05 is used for rejecting null hypothesis testing. No statistical methods were used to predetermine sample size randomization nor blinding was applied.

#### Area Under the Receiver Operating Characteristics (auROC)

We used auROC to classify neurons into different response classes to Ensure and water deliveries during slow feeding (Botta et al. 2020; Cohen et al. 2012). In individual neurons, we compared the distribution of raw Ca^2+^ amplitudes during baseline activity (−1 to 0 s before pump activation) to the distribution of raw Ca^2+^ amplitudes in individual 100-ms bins across trials. To produce the bin-specific ROC curves, we moved a criterion from the minimal to the maximal Ca^2+^ response we found in the neuron’s baseline activity distribution and the given 100 ms bin distribution. We then plotted the probability that Ca^2+^ signals in the given 100ms bin distribution were larger than the criterion against the probability that the Ca^2+^ signals in the baseline distribution were larger than the criterion. The auROC for each bin was then calculated using the auc function (sklearn.metrics.auc), resulting in auROC values between 0-1 and 0.5 means not different from the baseline. The post-stimulus auROC values from each time bin were compared to the baseline auROC values. Significance was established if at least four consecutive post-stimulus bin values between 0-2 s were greater than 2 S.D. of the pre-stimulus baseline values (food- or water-activated neurons).

#### Effect size calculation

To calculate the effect size of binge feeding-induced suppression in each neuron class, we performed a bootstrap-coupled estimation (DABEST, Python). To obtain the distribution of the mean difference between the two conditions, we re-sampled the mean Ca^2+^ activity of two conditions (eg. slow feeding and binge feeding) 5000 times (bootstrapping distribution, represented as the violin plot in Fig. 2e, 3m, and Fig S8a). The distribution was then normalized by the pooled standard derivation of both conditions to convert it to Cohen’s d using the Dabest.cohens_d (DABEST). P values were computed with the Dabest.PermutationTest (DABEST).

#### Q values calculation

To estimate the differences between 2 neuronal time series along the time axis, we first calculated the P values of each time bin by performing the unpaired Student’s t-test (scipy.stats.ttest_ind) and then applied false discovery rate correction (statsmodels.stats.multitest.fdrcorrection) to obtain the adjusted P values, which are the Q values.

#### Linear Mixed Models

Linear mixed models were used to estimate contributions of predictors (eg. level of excitation/suppression in neuronal activities upon feeding, or interaction of optogenetic actuators and light stimulation) to outcome (eg. food consumption or feeding bout duration) while allowing different intercepts for individual mice (lmer, R). To calculate the contribution of a given predictor, we built a full model with all predictors and a reduced model that lacks the given predictor. We then compared these two models with anova function (R) to calculate the P value of the given predictor. Representative models are structured as following:

### Full model

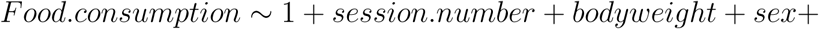

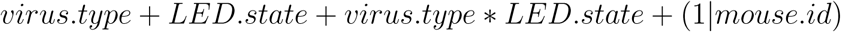

### Reduced model

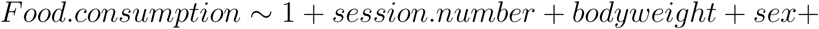

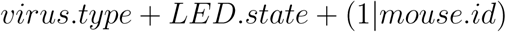

### Histology and imaging

Mice were anesthetized with (100 mg/Kg) Ketamine and (15 mg/Kg) Xylazine and perfused with 0.1M PBS and then 4% paraformaldehyde (PFA). Mice brains were harvested and stored in 4% PFA at 4°C overnight and transferred to 0.1M PBS for long-term storage. Brains were embedded in 4% agar-agar and sliced at 100-150 µm thickness with a vibratome. Brain slices were mounted on glass slides and were imaged by an epifluorescence microscope (Leica DMi8) or a confocal microscope (Leica SP5). Acquired images were then aligned to the mouse brain atlas (Paxinos and Keith B. J. Franklin 2007) for registration of the location of GRIN lens-prism or fiber optic cannula implants.

### Serial 2-Photon tomography (Brainsaw)

A subset of fixed mice brains was sliced and imaged by serial 2P tomography, where whole forebrain structures can be imaged at cellular resolutions (Ragan et al. 2012). We modified a custom-made 2P microscope (COSYS, UK) to operate with BakingTray (ScanImage & BakingTray, MATLAB). Obtained images were stitched (StitchIt, MATLAB) and reconstructed into 3D brain models. Image stacks were then registered to the Allen mouse brain atlas (BrainReg, Python) and visualized with napari (Python).

## SUPPLEMENTARY FIGURES

**Figure S1.**
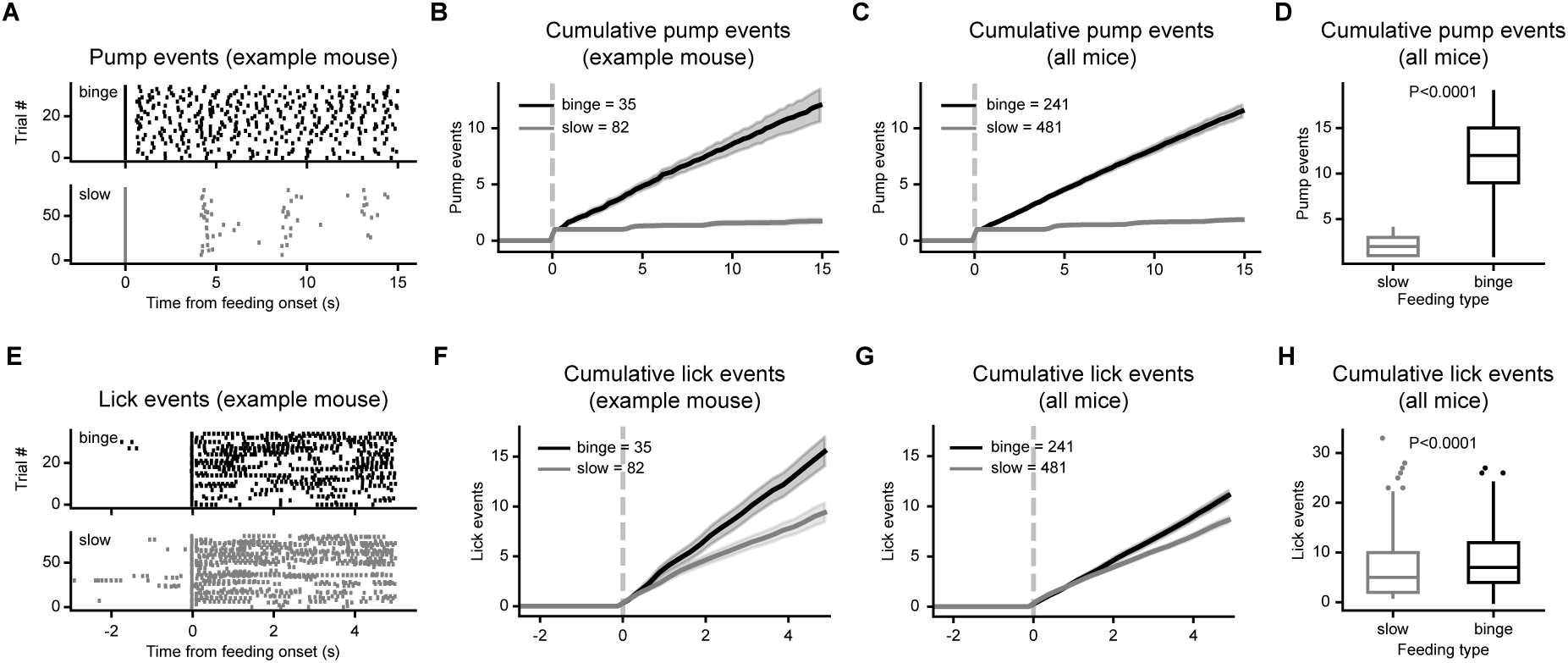
Behavioral differences between slow feeding and binge feeding. (**A**) Pump delivery events for slow feeding and binge feeding from an example mouse. (**B**) Cumulative pump events for slow feeding and binge feeding from an example mouse (same data as in **A**). (**C**) Same as in (**B**) but for all aPC CaMK2^+^ mice (n= 481 for slow feeding trials and 241 trials for binge feeding trials from 8 mice). (**D**) Quantification of cumulative pump events within 15 s after feeding bout onset. Unpaired Student’s t-test (n is the same as in **C**). (**E**) Lick events for slow feeding and binge feeding from an example mouse. (**F**) Cumulative lick events for slow feeding and binge feeding from an example mouse (same data as in **E**). (**G**) Same as in (**F**) but for all aPC CaMK2^+^ mice (n is the same as in **C**). (**H**) Quantification of cumulative lick events in 4 s after the onset of feeding bouts. Unpaired Student’s t-test (n is the same as in **C**). For (**B**) (**C**) (**F**) (**G**) data are shown as mean ± s.e.m. For the box plot in (**D**) (**H**) the center line shows the median, the box limits show the quartiles, the whiskers show 1.5x the interquartile range, and the points show the outliers.

**Figure S2.**
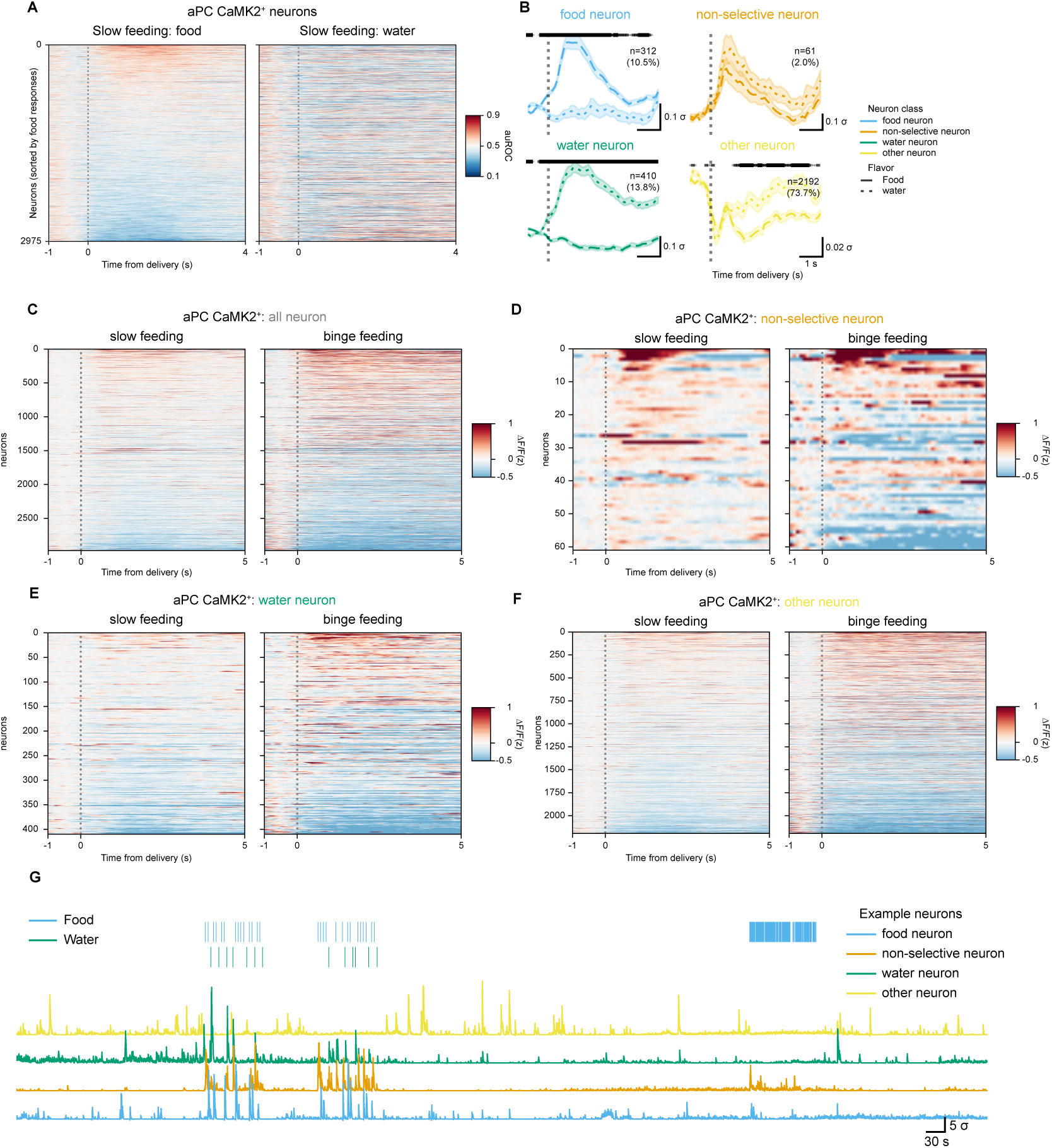
aPC CaMK2^+^ neuronal responses upon slow feeding and binge feeding. (**A**) Area Under the Receiver Operating Characteristics (auROC) curve of aPC CaMK2^+^ neurons during slow feeding (see **Methods**, n is 2975 cells from 8 mice). (**B**) Slow feeding responses of aPC CaMK2^+^ neuronal subclasses for food and water deliveries (n=312 food neurons, 61 non-selective neurons, 410 water neurons, 2192 other neurons from 8 mice). The shaded line above denotes the adjusted P-values (Q-values) of each time point, with different line widths representing different values (from thin to thick: q <0.05, <0.01, <0.001). (**C**) Responses of individual aPC CaMK2^+^ neurons upon slow feeding and binge feeding (n is the same as in **A**). (**D**) Same as in (**C**) but with non-selective consumption neurons (n=61 cells from 8 mice). (**E**) Same as in (**C**) but with water-activated neurons (n=410 cells from 8 mice). (**F**) Same as in (**C**) but with non-responding neurons (n=2192 cells from 8 mice). (**G**) Example traces of aPC CaMK2^+^ neurons from a recording session. For (**B**) data are shown as mean ± s.e.m.

**Figure S3.**
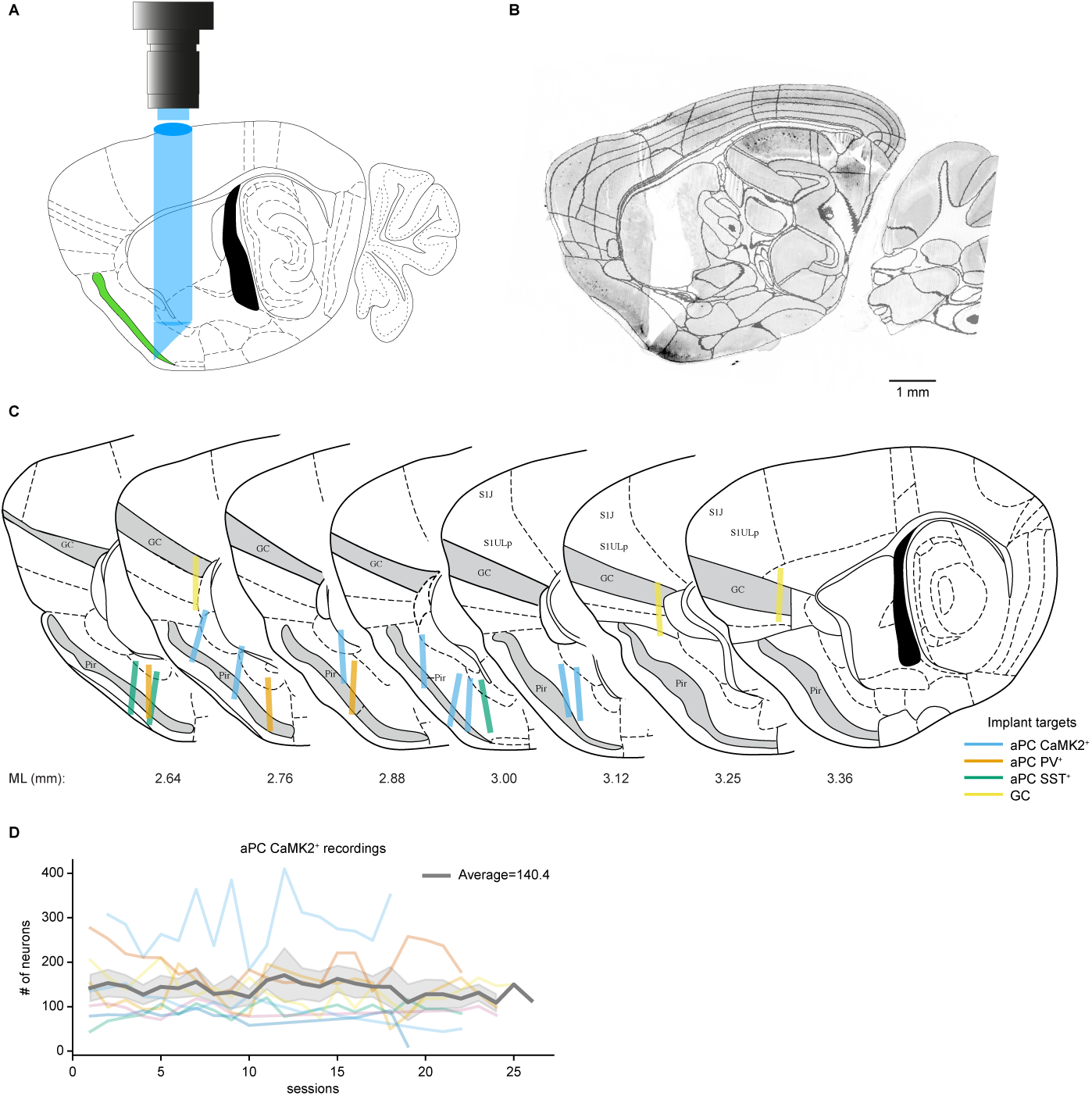
GRIN lens-Prism Implant coordinates and extracted cell numbers throughout experimental sessions. (**A**) Schematic of implant coordination in the aPC. (**B**) GRIN lens-Prism path and GCaMP6f expression from an example mouse. (**C**) Reconstructed GRIN len-Prism coordinates in different cell types in the aPC (Fig.1, Fig.3) and GC (Fig.2). (**D**) Numbers of aPC CaMK2^+^ neurons extracted with *CNMFe* on each recording session (n= 8 mice). For (**D**) data are shown as mean ± s.e.m.

**Figure S4.**
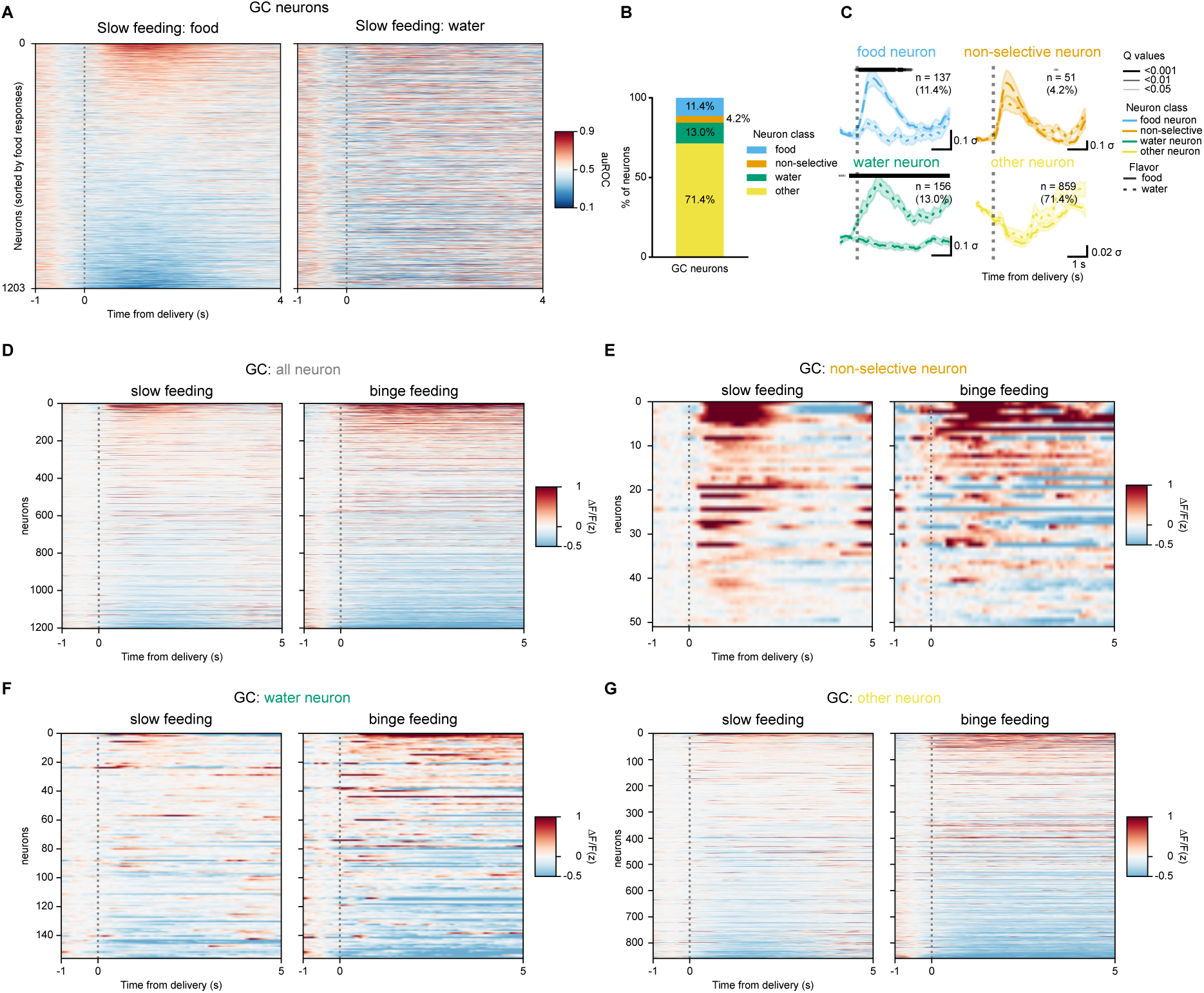
GC neuronal responses upon slow feeding and binge feeding. (**A**) Area Under the Receiver Operating Characteristics (auROC) curve of GC neurons during slow feeding (see **Methods**). (**B**) Proportion of GC subclasses (n=1203 all neurons, 137 food neurons, 51 non-selective neurons, 156 water neurons, and 859 other neurons from 3 mice). (**C**) Food responses of subclasses of GC neurons during slow feeding for food and water deliveries (n is the same as in **B**). The shaded line above denotes the adjusted P-values (Q-values) of each time point, with different line widths representing different values (from thin to thick: q <0.05, <0.01, <0.001). (**D**) Responses of all GC neurons upon slow feeding and binge feeding (n=1203 cells from 3 mice). (**E**) Same as in (**D**) but with non-selective consumption neurons (n=51 cells from 3 mice). (**F**) Same as in (**D**) but with water-activated neurons (n=156 cells from 3 mice). (**G**) Same as in (**D**) but with non-responding neurons (n=859 cells from 3 mice). For (**C**) data are shown as mean ± s.e.m.

**Figure S5.**
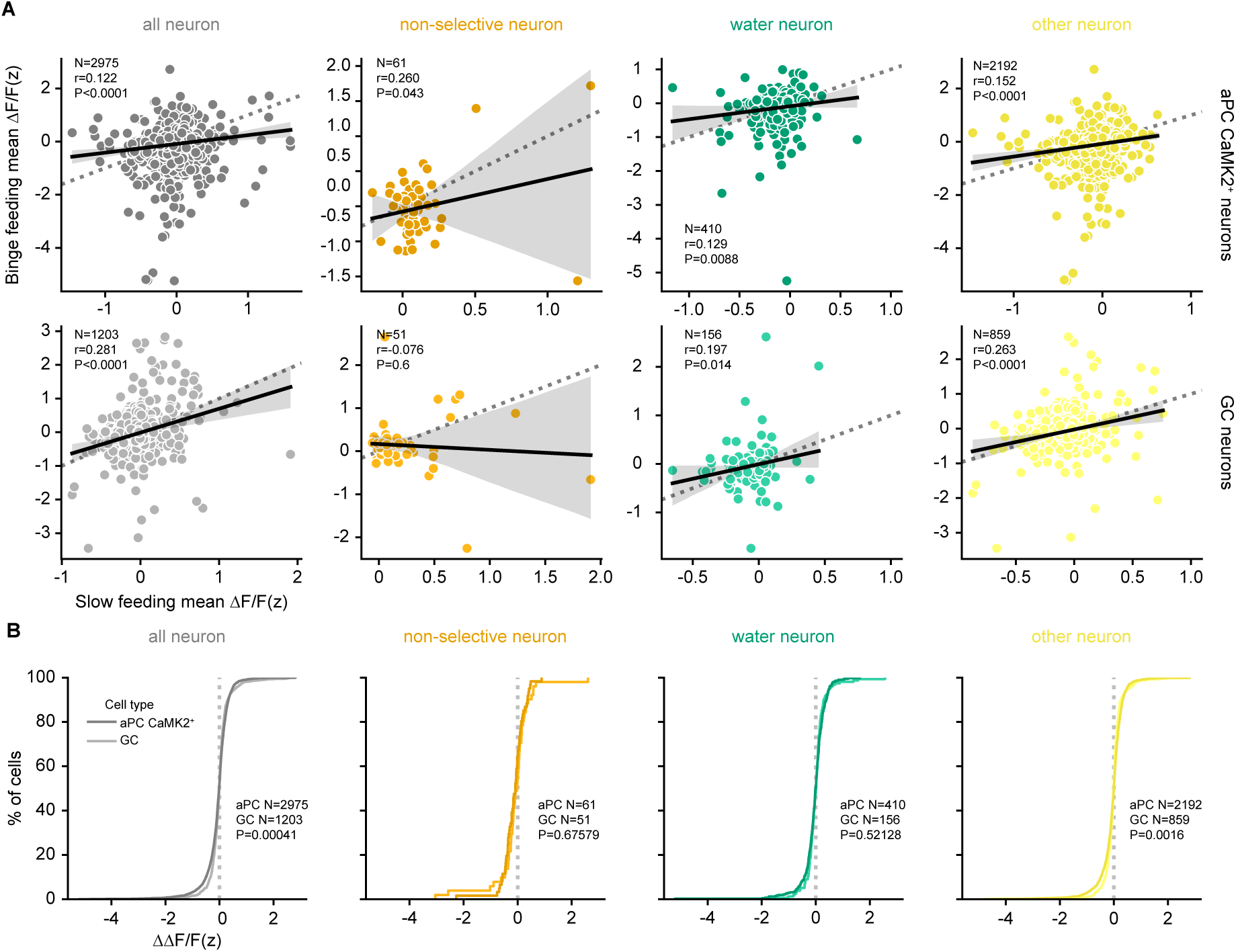
Distinct binge feeding-induced modulation in the aPC and the GC. (**A**) Cell-wise comparison of neuronal responses upon slow feeding and binge feeding (For aPC CaMK2^+^ neurons, n=2975 for all neurons, 61 for non-selective neurons, 410 for water neurons, 2192 for other neurons from 8 mice. For GC neurons, n=1203 for all neurons, 51 for non-selective neurons, 156 for water neurons, and 859 for other neurons from 3 mice). (**B**) Cumulative distribution of the difference for each cell upon slow and binge feeding (binge feeding - slow feeding, ΔΔ F/F(z), n is the same as in **A**. In (**A**) r and P represent the correlation coefficient and P-value of Pearson’s r.

**Figure S6.**
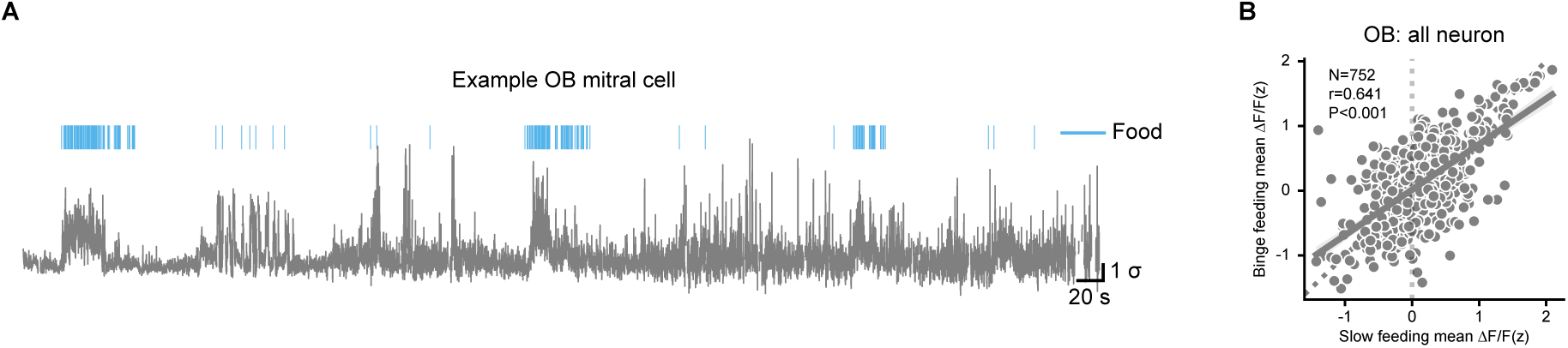
OB mitral cell example trace and cell-wise comparison. (**A**) An example trace from an OB mitral cell upon slow feeding and binge feeding. (**B**) Cell-wise comparison of neuronal responses upon slow feeding and binge feeding (n=752 cells from 4 mice). In (**B**) r and P represent the correlation coefficient and P-value of Pearson’s r.

**Figure S7.**
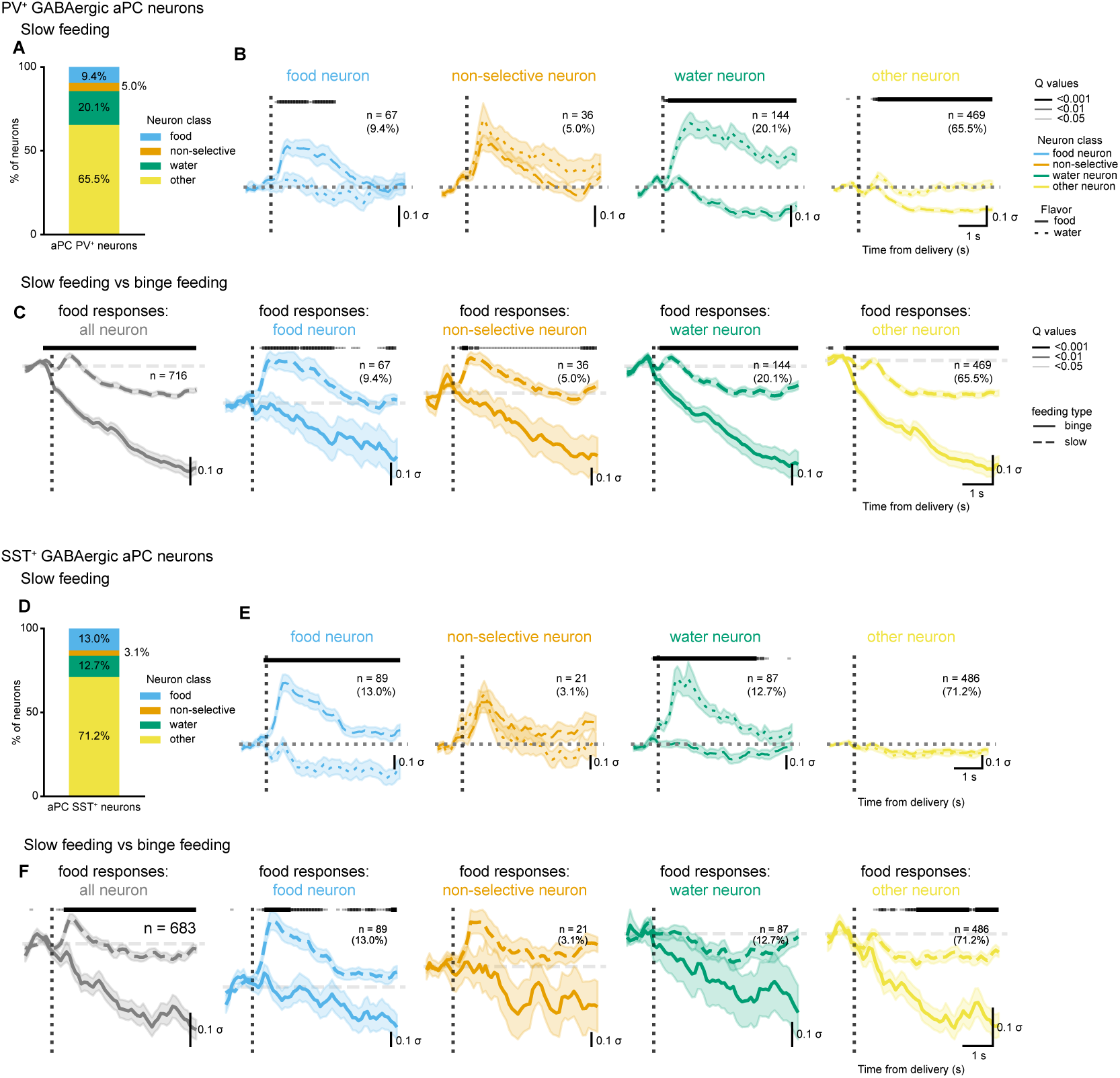
aPC GABAergic neuronal responses to slow feeding and binge feeding. (**A**) Proportion of aPC PV^+^ subclasses (n=67 cells for food neurons, 36 cells for non-selective neurons, 144 cells for water neurons, 469 cells for other neurons from 3 mice). (**B**) Responses of subclasses of aPC PV^+^ neurons during slow feeding for food and water deliveries (n is the same as in **a**, except n= 716 cells for all neurons). (**C**) Food responses aPC PV^+^ neuron subclasses during slow feeding and binge feeding (n is the same as in **B**). (**D**) Same as in (**A**) but for aPC SST^+^neurons (n=89 cells for food neurons, 21 cells for non-selective neurons, 87 cells for water neurons, 486 cells for other neurons from 3 mice). (**E**) Same as in (**B**) but for aPC SST^+^neurons (n is the same as in **D**). (**F**) Same as in (**C**) but for aPC SST^+^neurons (n is the same as in (**E**) except n=683 cells in all neurons). The shaded line above denotes the adjusted P-values (Q-values) of each time point, with different line widths representing different values (from thin to thick: Q <0.05, <0.01, <0.001). For (**B**) (**C**) (**E**) (**F**) data are shown as mean ± s.e.m.

**Figure S8.**
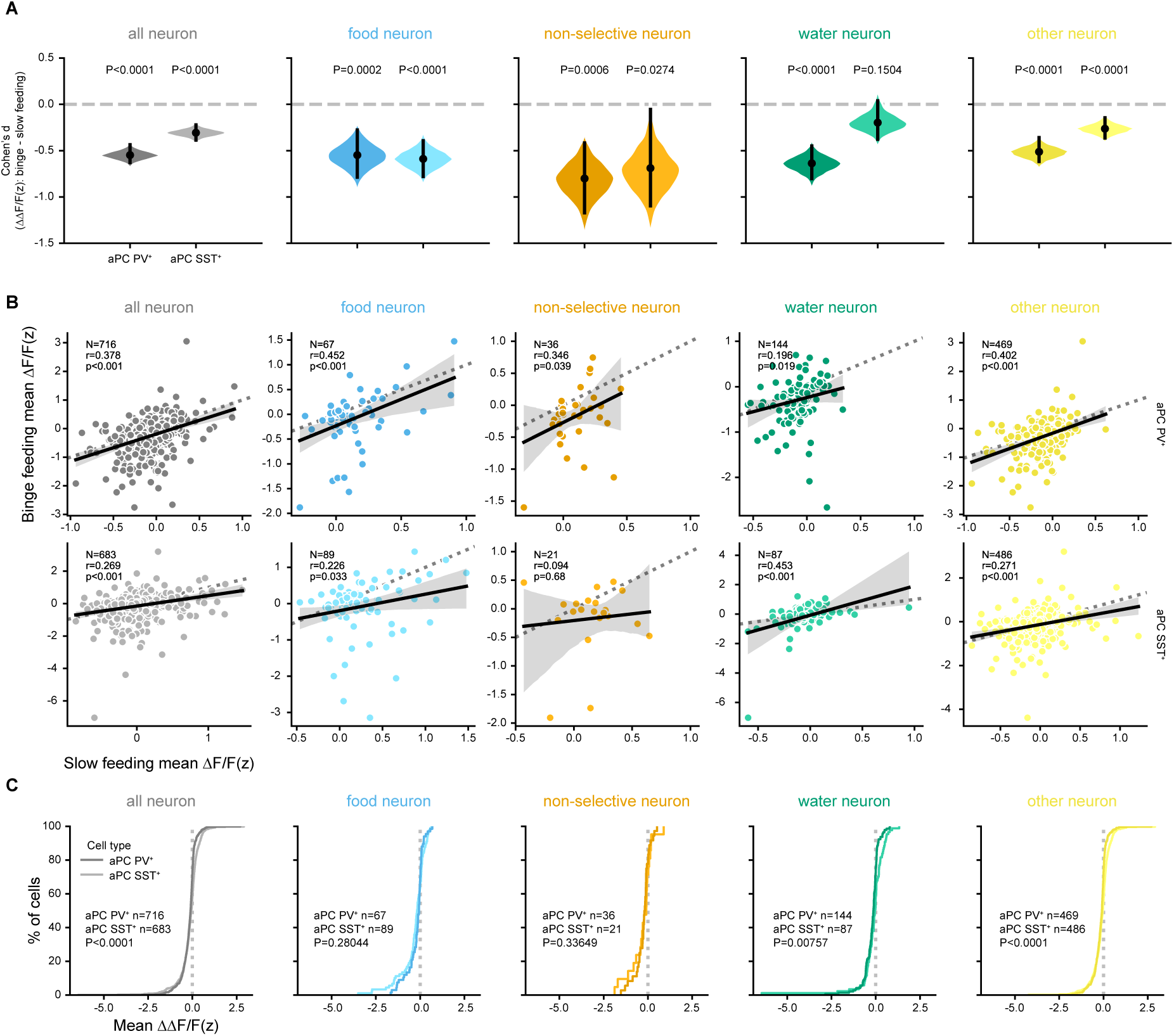
Cell-wise comparison of aPC GABAergic neuronal responses to slow feeding and binge feeding. (**A**) Estimated effect size of binge feeding-induced suppression in aPC PV^+^ and aPC SST^+^ subclasses (For aPC PV^+^ neurons, n=716 cells for all neurons, 67 cells for food neurons, 36 cells for non-selective neurons, 144 cells for water neurons, 469 cells for other neurons from 3 mice. For aPC SST^+^ neurons, n=683 cells for all neurons, n=89 cells for food neurons, 21 cells for non-selective neurons, 87 cells for water neurons, 486 cells for other neurons from 3 mice). (**B**) Cell-wise comparison of neuronal responses upon slow feeding and binge feeding in aPC PV^+^ and aPC SST^+^ subclasses (n is the same as in **A**). (**C**) Cumulative distribution of the difference for each cell upon slow and binge feeding (binge feeding - slow feeding, ΔΔ F/F(z)) in aPC PV^+^ and aPC SST^+^ subclasses (n is the same as in **A**). For (**B**) r and P represent the correlation coefficient and P-value of Pearson’s r.

**Figure S9.**
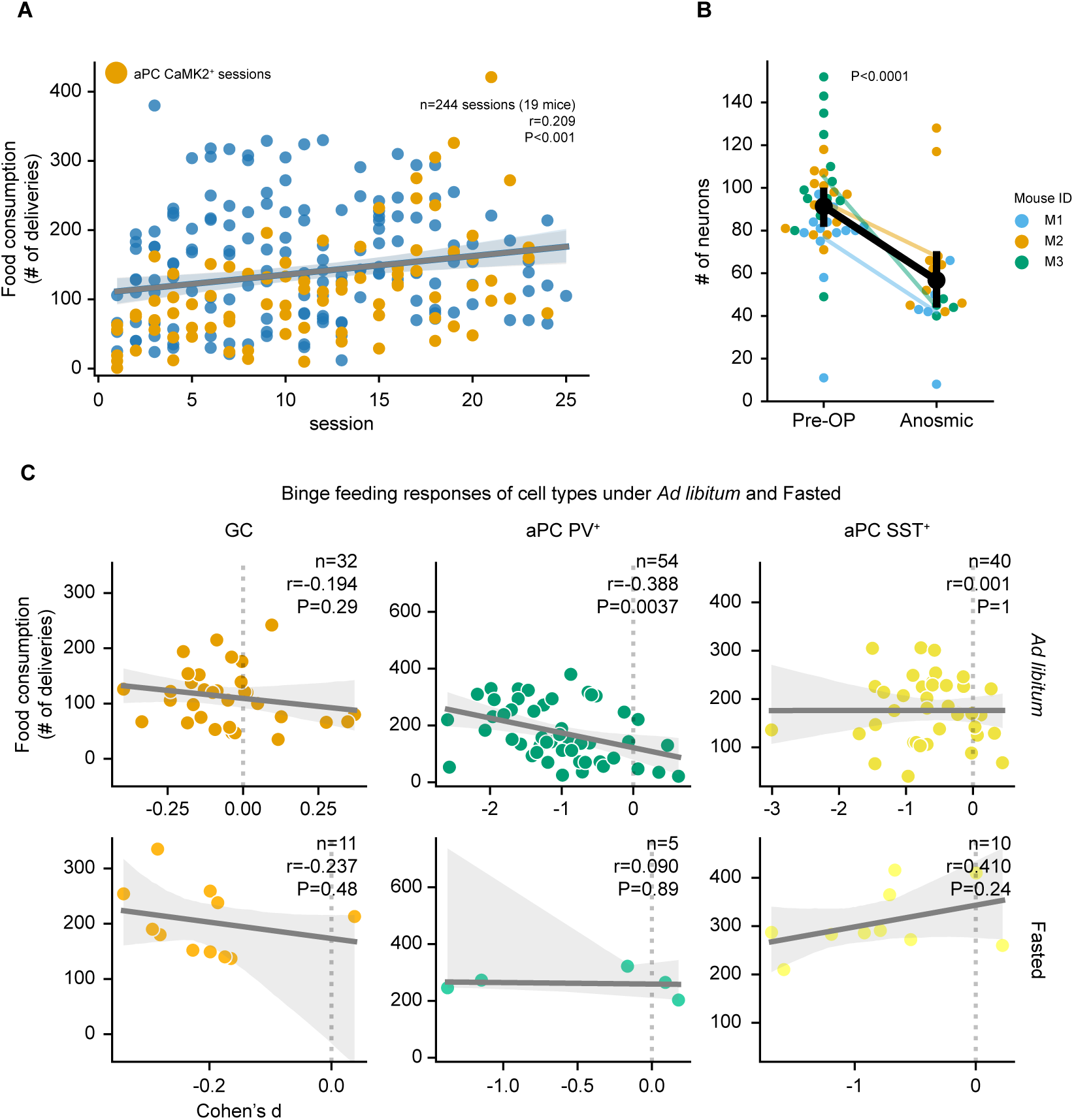
Effects of temporal progression, anosmia, and fasting. (**A**) Food consumption (number of food deliveries) on subsequent recording sessions (n=244 sessions from 19 mice). Orange dots are mice with recordings in aPC CaMK2^+^ cells (n=84 sessions from 8 mice), blue dots represent all other mice. (**B**) Number of extracted cells in aPC CaMK2^+^ mice before and after Triton-X100 application (n=21 Pre-OP sessions and 15 anosmic sessions from the same 3 mice). Colors represent recordings from each mouse. **c**, Neuronal responses to binge feeding in the GC, aPC PV^+^, and aPC SST^+^under *ad libitum* (upper row) and fasting (lower row) conditions plotted against consumption (For GC, n=32 *ad libitum* sessions and 11 fasted sessions from 3 mice. For aPC PV^+^, n=54 *ad libitum* sessions and 11 fasted sessions from 3 mice. For aPC SST^+^, n=40 *ad libitum* sessions and 10 fasted sessions from 3 mice.). For (**B**) data are shown as mean ± s.e.m. For (**A**) (**C**) r and P represent the correlation coefficient and P-value of Pearson’s r.

**Figure S10.**
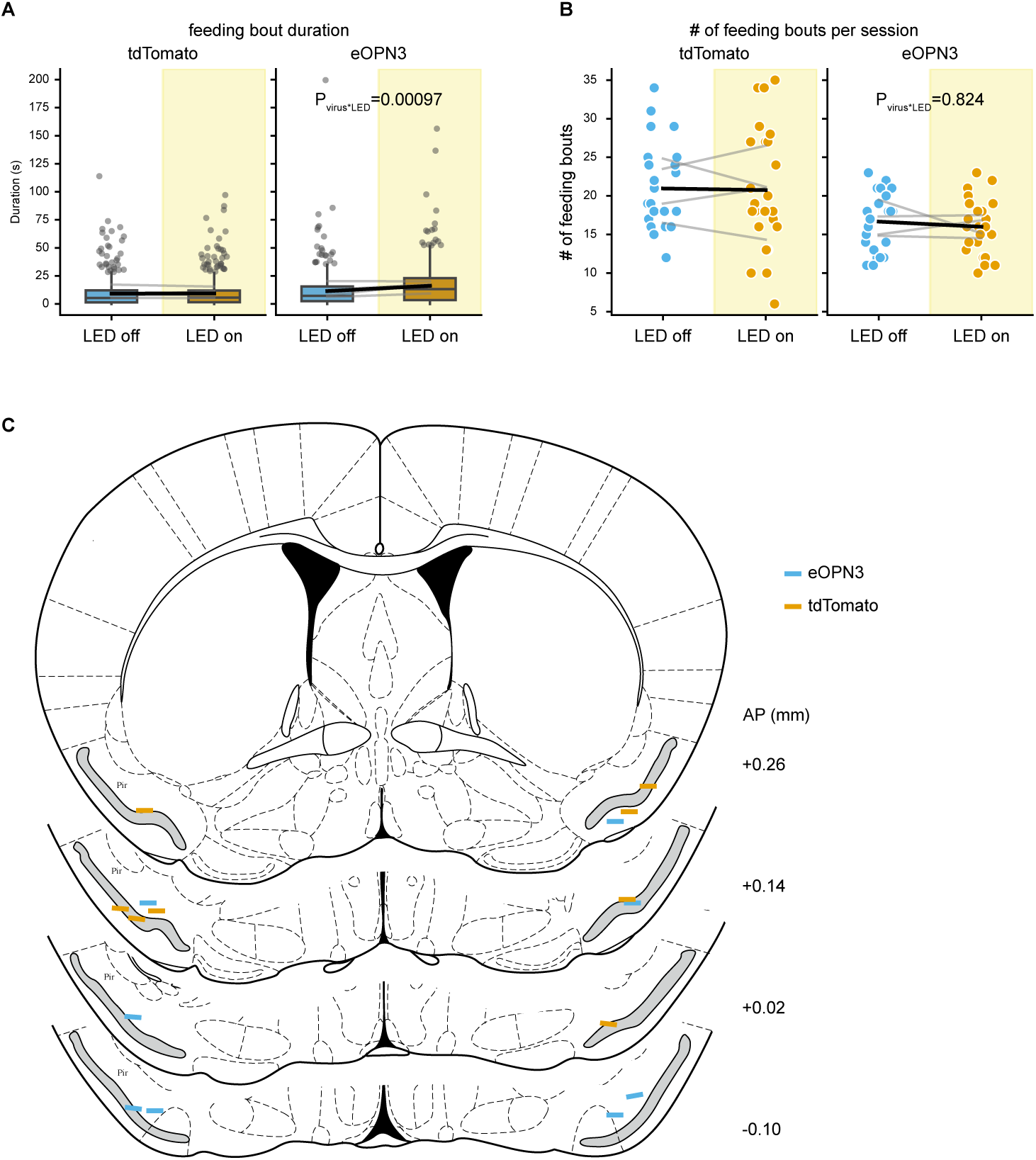
Effects of optogenetic suppression of aPC and implant coordinates. (**A**) Effects of light stimulation on the duration of individual feeding bouts in tdTomato- and eOPN3-expressing mice (n=569 feeding bouts in LED off sessions and 562 feeding bouts in LED on sessions from 4 tdTomato mice. n=431 feeding bouts in LED off sessions and 425 feeding bouts in LED on sessions from 4 eOPN3 mice). (**B**) Effects of light stimulation on the number of feeding bouts per experimental session in tdTomato- and eOPN3-expressing mice (n=24 LED off sessions and 24 LED on sessions from 4 tdTomato mice and n=24 LED off sessions and 24 LED on sessions from 4 eOPN3 mice). Grey lines denote the data from the same mice, and the black line denotes the overall mean of the data. The p values are calculated from a linear mixed model. (**C**) Implant coordinates of optical fibers. For the box plot in a and b, the center line shows the median, the box limits show the quartiles, the whiskers show 1.5x the interquartile range, and the points show the outliers.

**Figure S11.**
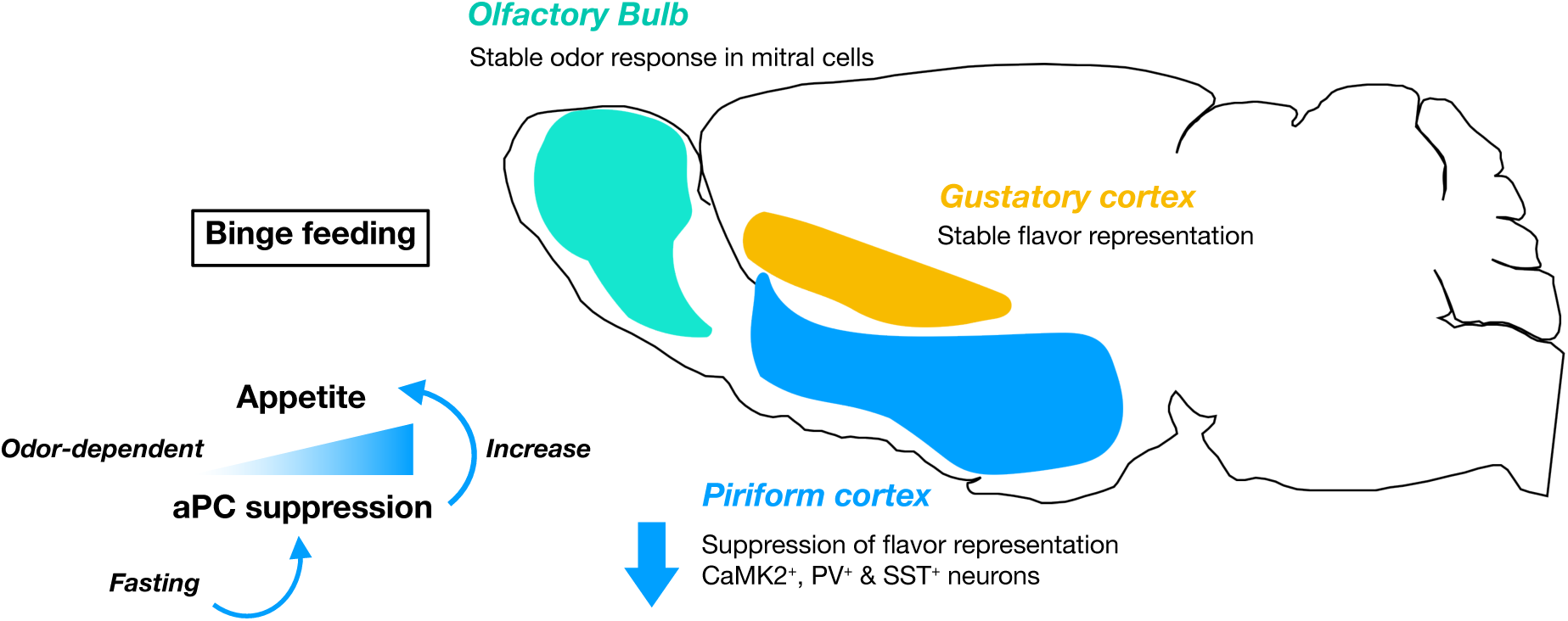
Graphic summary. In this study, we found a feeding rate-dependent suppression of the olfactory flavor representation (Fig. 1 and Fig. 3) whereas the gustatory flavor representation is not affected by the feeding rate (Fig. 2). Olfactory inputs from the olfactory bulb remain stable across feeding rates (Fig. 3). We found the magnitude of binge feeding-induced aPC suppression correlates with appetite and the correlation depends on olfactory perception and metabolic state (Fig. 4). We further showed optogenetically suppression in the aPC upon feeding promotes appetite (Fig. 5).

## SUPPLEMENTARY VIDEOS

**Video. S1:**
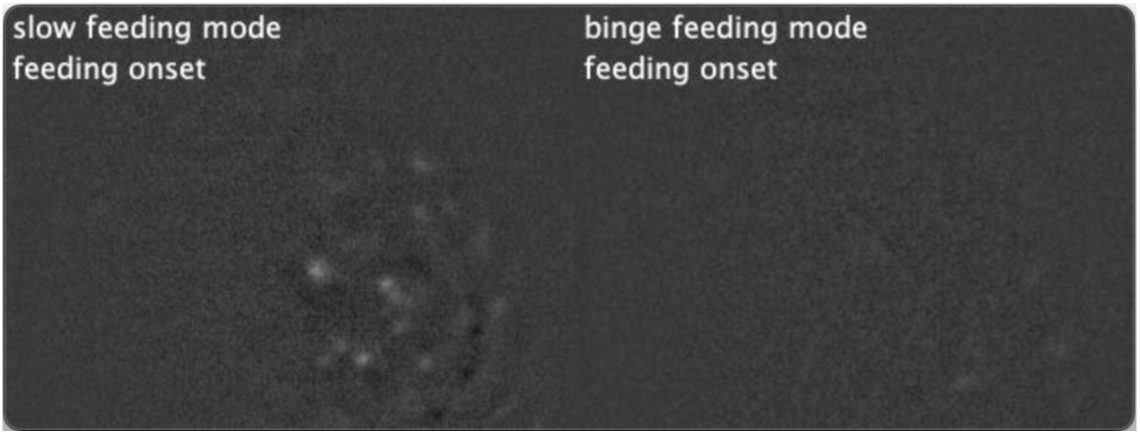
Example Ca^2+^ video during slow feeding and binge feeding. Left panel: Ca^2+^ transients of aPC CaMK2^+^ neurons during slow feeding. Right panel: Ca^2+^ transients of aPC CaMK2^+^ neurons during binge feeding from the same recording, the same mice.

## SUPPLEMENTARY TABLES

### Coordinations for injection and implantation

**Table S1:** Coordinates for viral injection and GRIN lens and prism implantation.

| Brain regions | Coordinates<br>(AP/ML/DV, mm) | Notes |
| --- | --- | --- |
| aPC | 0.32/-3.1/5.2<br>0.32/-2.7/5.5 | For viral injection<br>For implantation we used the bottom right corner of the prism of GRIN lens |
| GC | 0.26/-3.6/4.0<br>0.26/-3.1/4.1 | For viral injection<br>For implantation we used the bottom right corner of the prism of GRIN lens |
| OB | 4.3-4.6/±0.6 /0.3 | For viral injection |

### Virus and construct

**Table S2:** Virus and construct table.

| **Reagent or resource** | **Source** | **Identifier (RRID)** |
| --- | --- | --- |
| AAV1<br>pAAV.Syn.Flex.GCaMP6f.WPRE.SV40 | Virus: Addgene | 100833-AAV1<br>RRID:Addgene_100833 |
| AAV8<br>pAAV-Syn-iCre-RFP | Plasmid: Charité viral core<br>Virus: Charité viral core | Charité viral core id: BA-48c |
| AAV1<br>pGP-AAV-syn-jGCaMP7s-WPRE | Virus: Addgene | 104487-AAV1<br>RRID:Addgene_104487 |
| AAV9<br>pAAV-hSyn-Flex-OPN3-mScarlet-minWPRE | Plasmid: Charité viral core<br>Virus: Charité viral core | Charité viral core id: BA-575b |
| AAV9<br>hSyn-flox-tdTomato | Plasmid: Charité viral core<br>Virus: Charité viral core | Charité viral core id: BA-234a |

### Mouse line

**Table S3:**
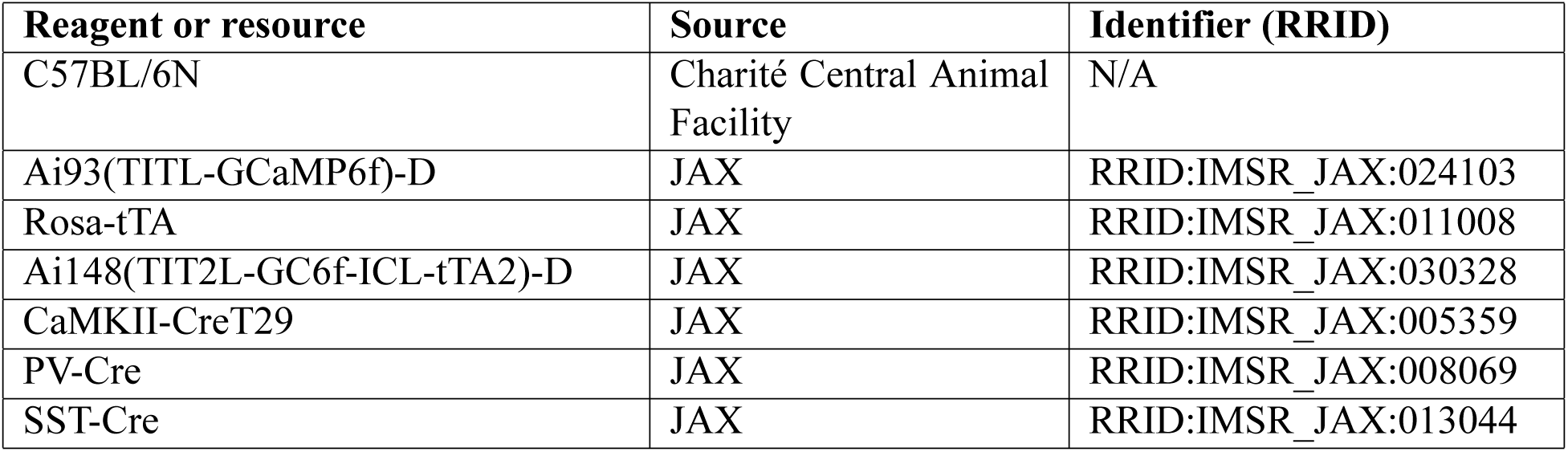
Mouse line table.

| Reagent or resource | Source | Identifier (RRID) |
| --- | --- | --- |
| C57BL/6N | Charité Central Animal Facility | N/A |
| Ai93(TITL-GCaMP6f)-D | JAX | RRID:IMSR_JAX:024103 |
| Rosa-tTA | JAX | RRID:IMSR_JAX:011008 |
| Ai148(TIT2L-GC6f-ICL-tTA2)-D | JAX | RRID:IMSR_JAX:030328 |
| CaMKII-CreT29 | JAX | RRID:IMSR_JAX:005359 |
| PV-Cre | JAX | RRID:IMSR_JAX:008069 |
| SST-Cre | JAX | RRID:IMSR_JAX:013044 |

### Software

**Table S4:**
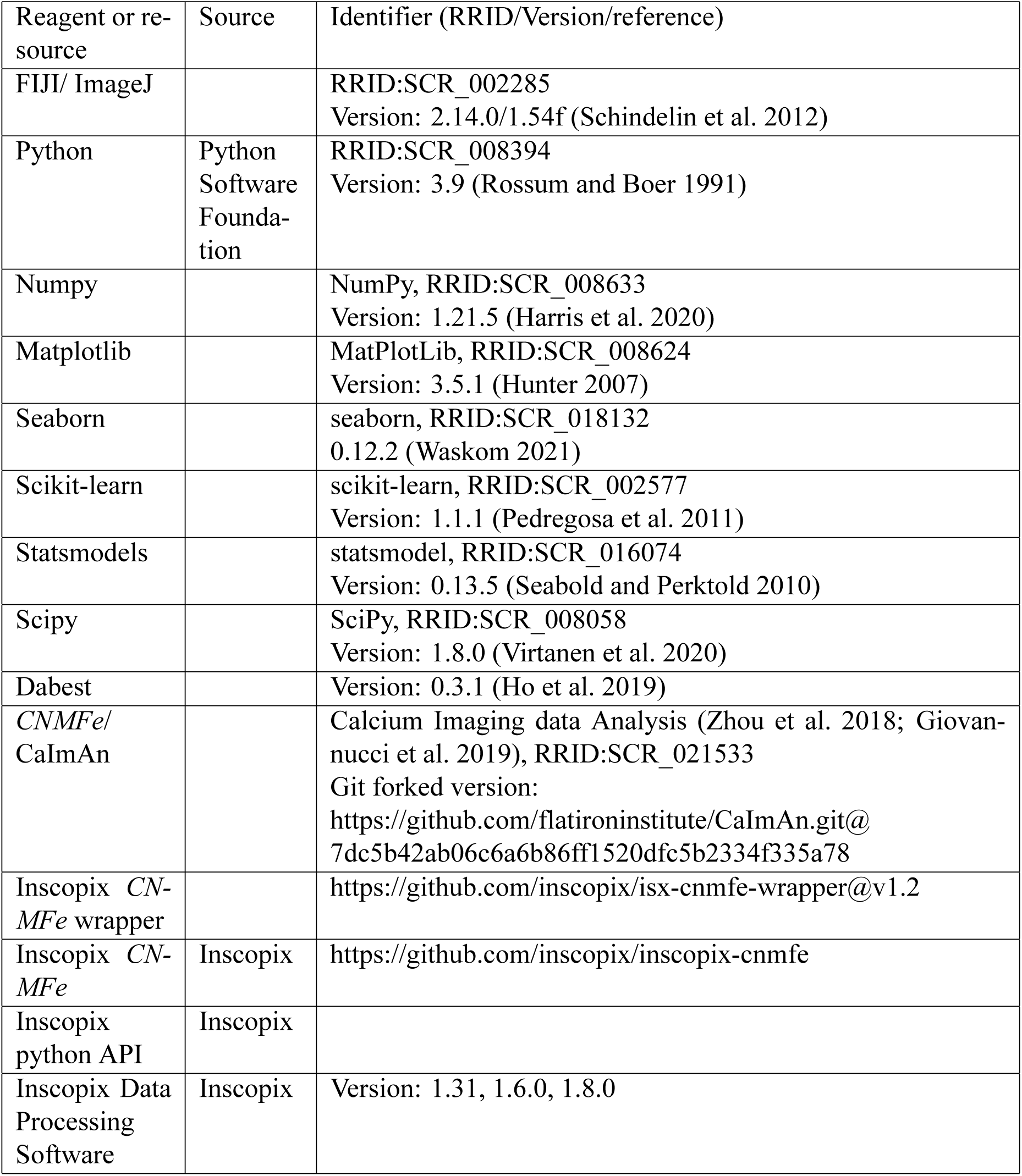
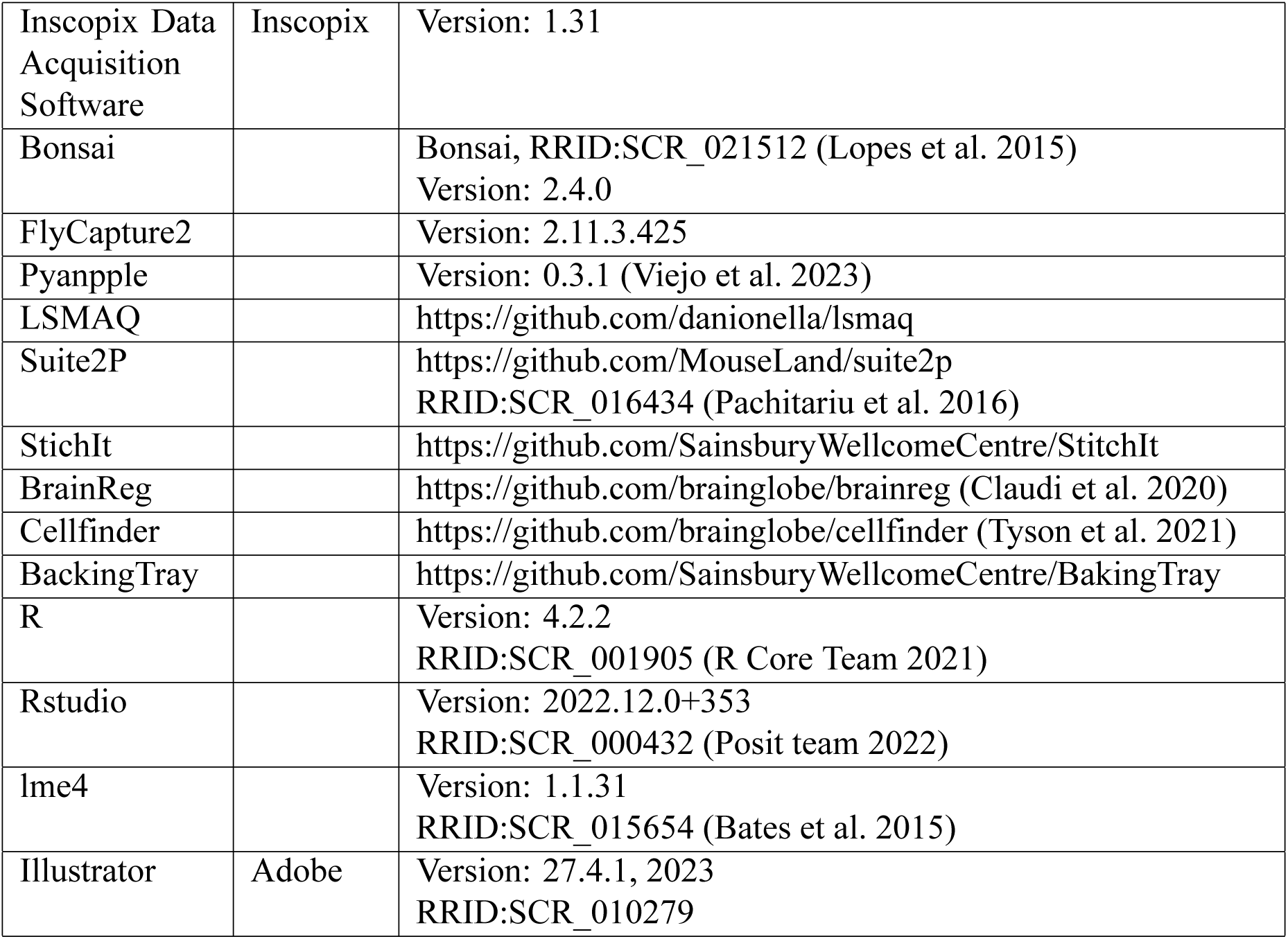
Software table.

### Statistical summary

**Table S5:**
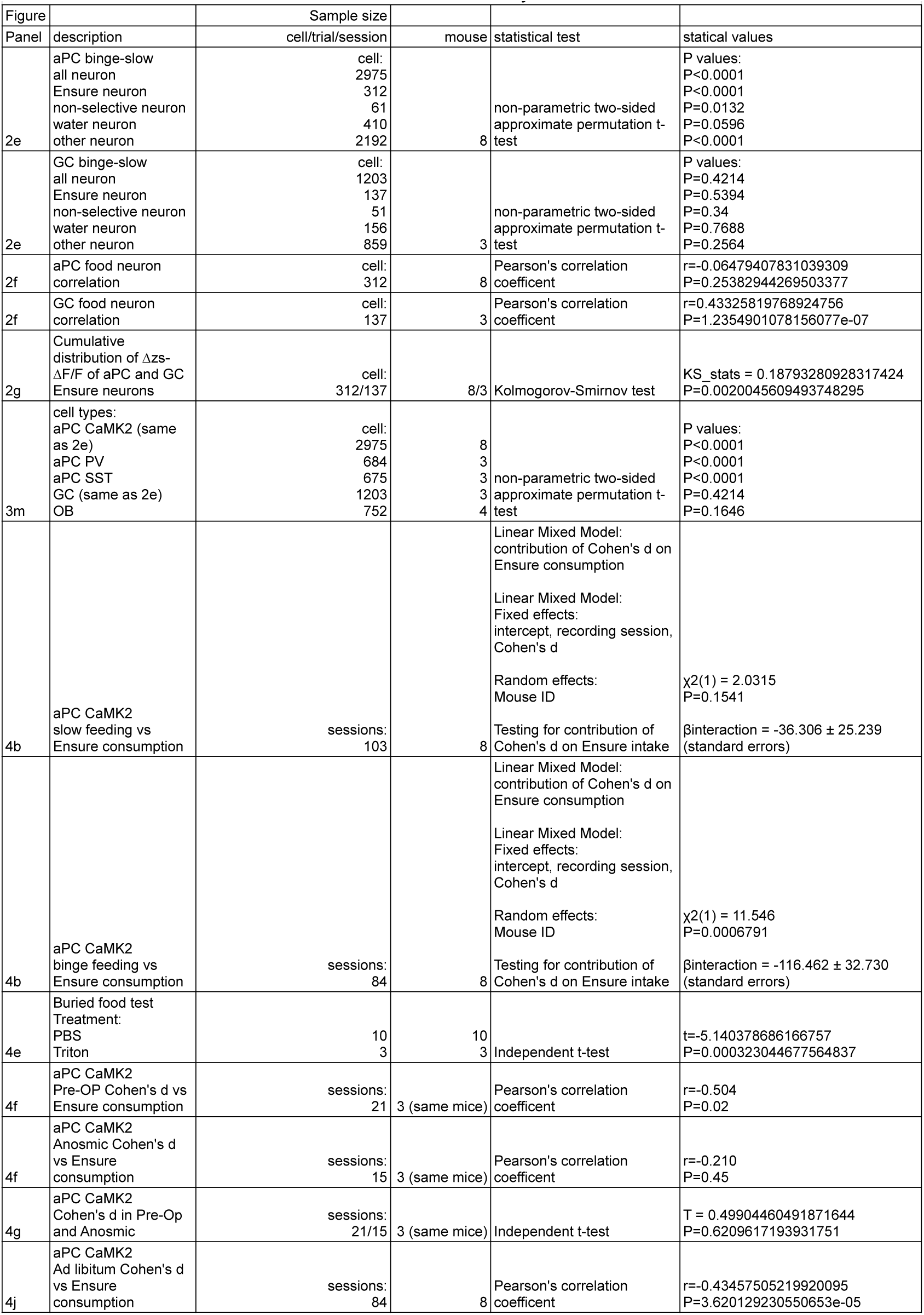
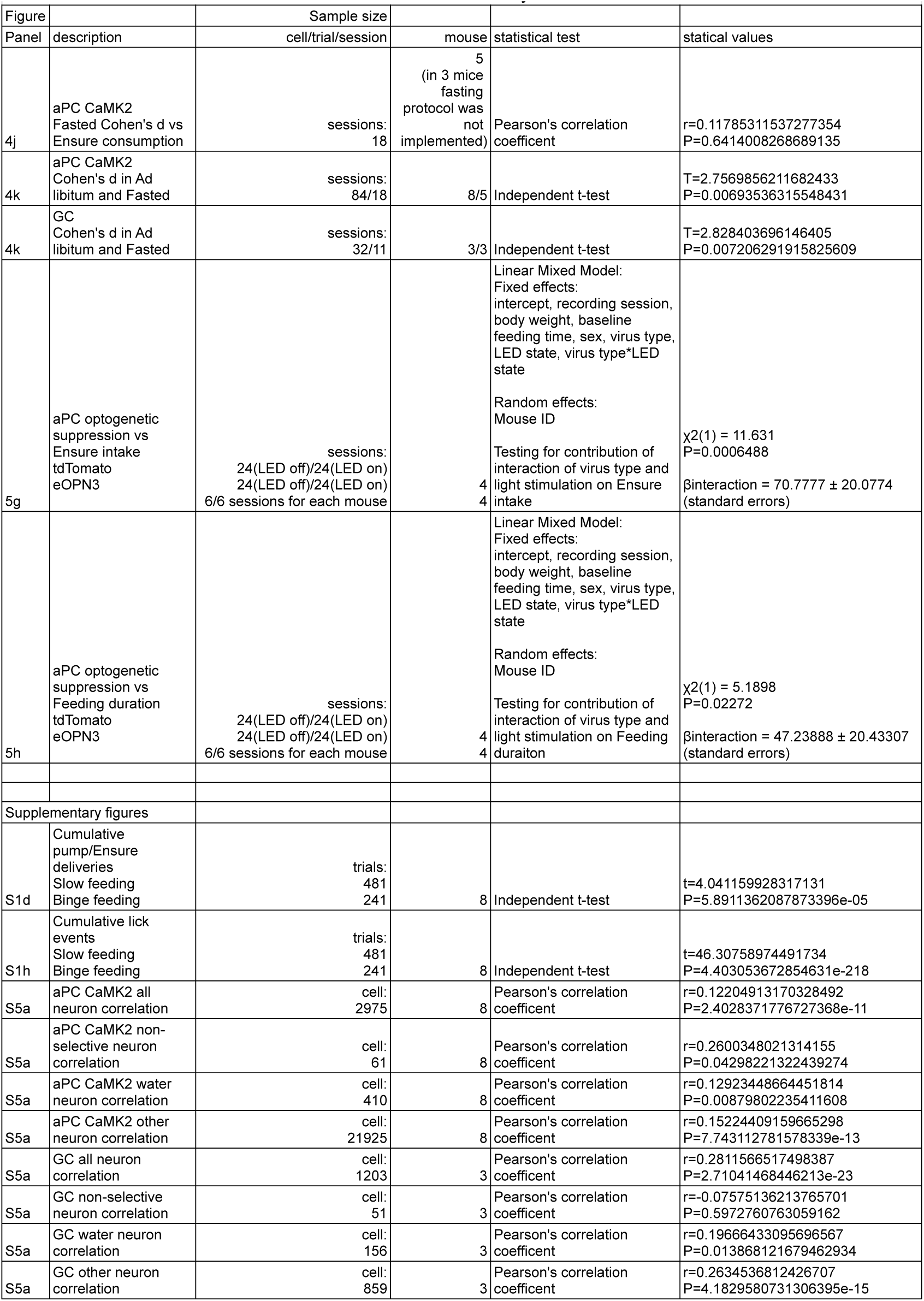
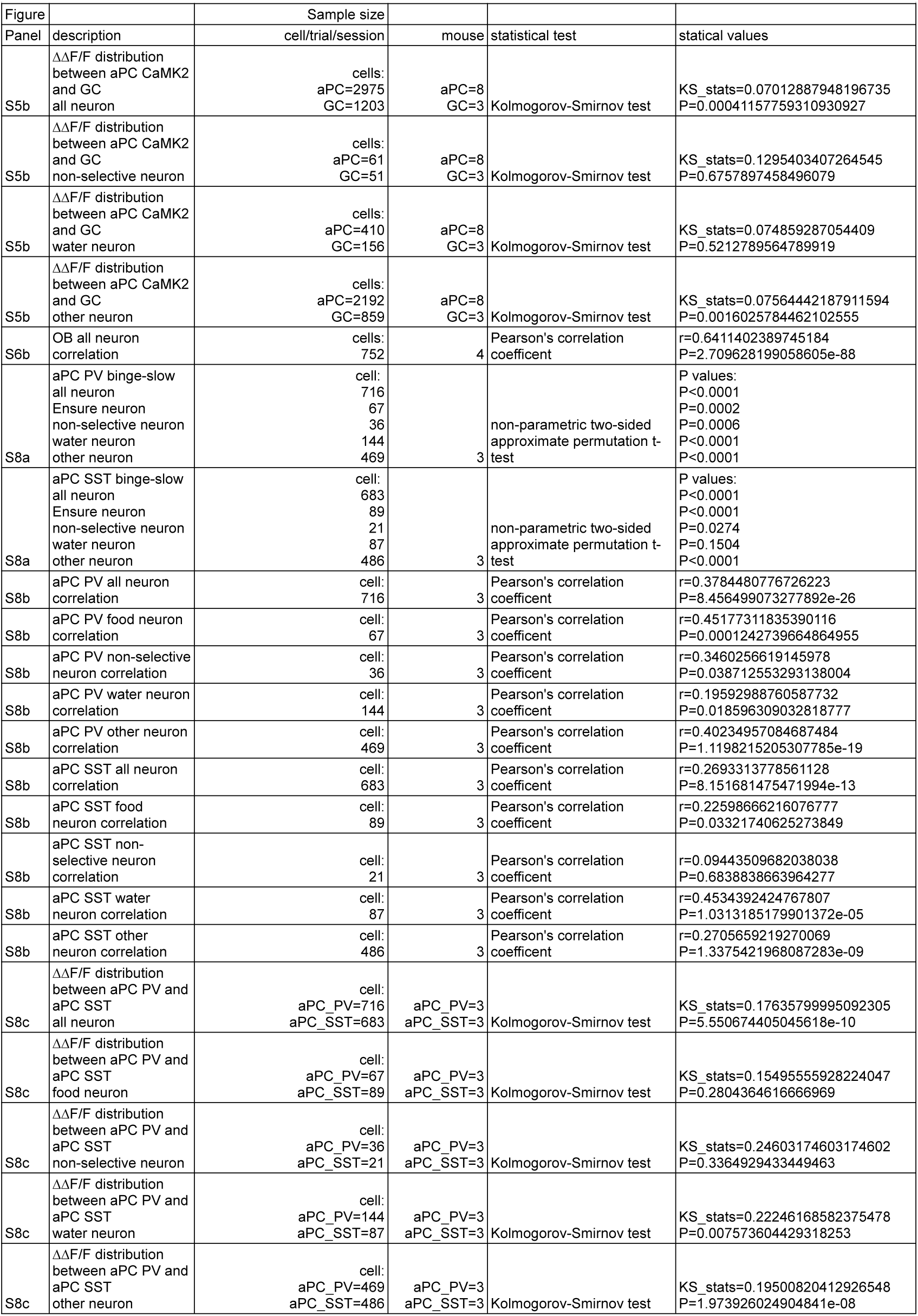
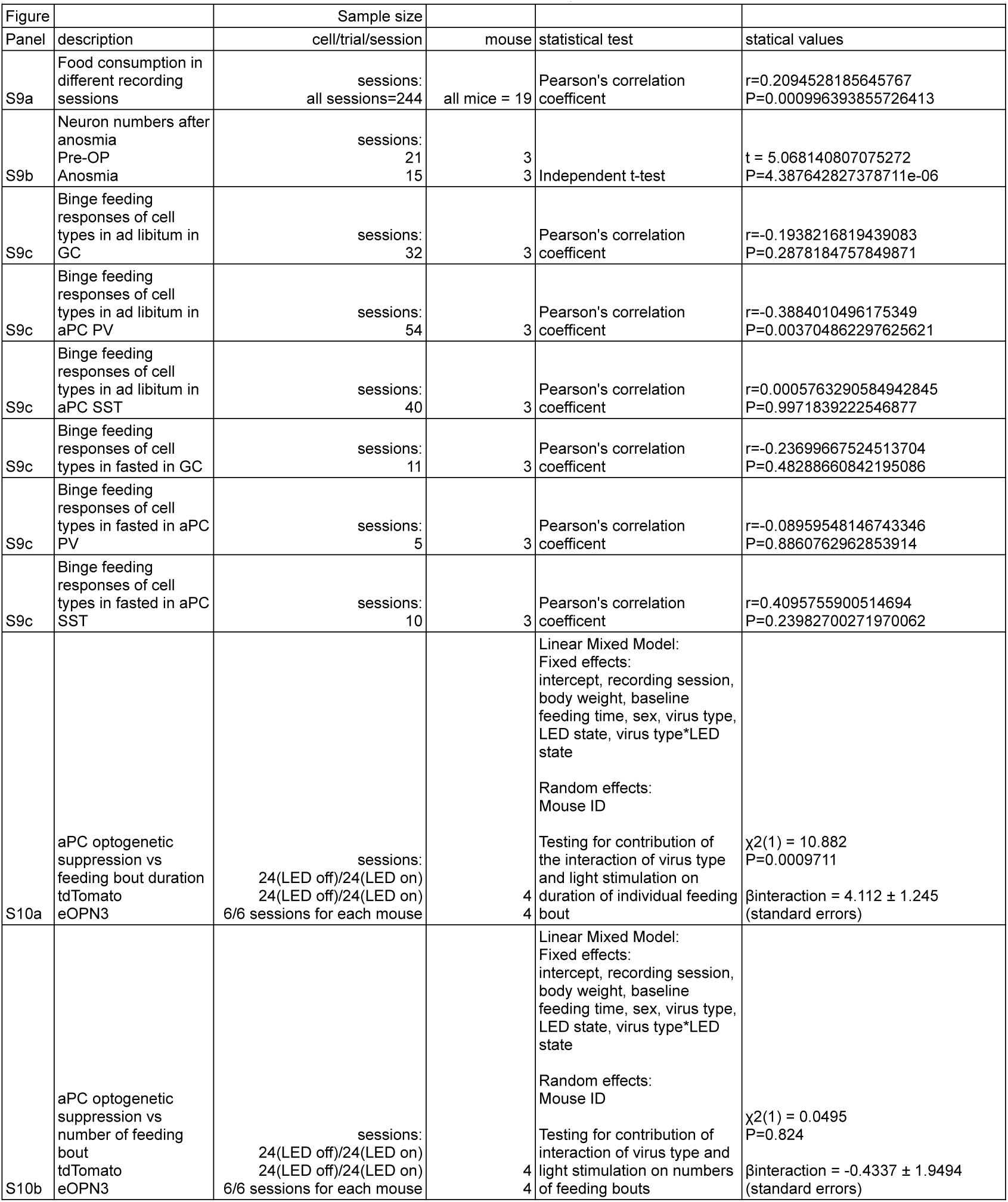
Statistical summary.

## Notes

### Summary of Updates

Update author names and affiliations, fix minor typos

